# *SLFN11* Loss-Induced Chemoresistance is Associated with Overexpression of Glycerophospholipid Biosynthesis in Ewing Sarcoma

**DOI:** 10.1101/2025.05.09.652501

**Authors:** Kasturee Chakraborty, Ritambhar Burman, Saharsh Satheesh, Matthew Kieffer, Chandni Karuhatty, Zuo-Fei Yuan, Haiyan Tan, Ankhbayar Lkhagva, Anthony A High, Xusheng Wang, Alaa Refaat, Weixing Zhang, Yaxu Wang, Yiping Fan, Madan M Babu, Anang Shelat, Elizabeth Stewart, Michael A Dyer, Puneet Bagga

## Abstract

Ewing sarcoma (EWS) is an aggressive cancer in adolescents and young adults with frequent relapse rates and poor outcomes in recurrent or metastatic cases. Schlafen family member 11 (*SLFN11*) gene is associated with the sensitivity to DNA-damaging agents (DDAs). The knockout of *SLFN11* is associated with acquired chemoresistance in both cell lines and preclinical models. Here, we aimed to elucidate the metabolic underpinnings of *SLFN11*-loss associated chemoresistance in patient derived cell lines of EWS. Our integrated transcriptomic and metabolomic analyses revealed downregulation of mitochondrial glycerol-3-phosphate dehydrogenase 2 (*GPD2*) gene, which was accompanied by the upregulation of glycerophospholipid (GPL) biosynthesis pathway. Further, therapeutic targeting of lipid synthesis with the glycerol-3-phosphate acyltransferase 1 *(GPAT1)* inhibitor (FSG67) enhanced the efficacy of the DDA (SN-38) in *SLFN11^−/−^* cells. These findings indicate that *SLFN11* loss-mediated chemoresistance can be targeted by blocking GPL biosynthesis in addition to DDA administration.

## Introduction

Ewing sarcoma (EWS) is an aggressive malignancy affecting adolescents and young adults, with high rates of relapse despite multimodal treatments.^1–3^ Both localized and metastatic tumors often recur, highlighting the need for novel therapeutic strategies and a better understanding of treatment resistance mechanisms.^3–7^ A hallmark of EWS tumors is a chromosomal translocation between the *EWSR1* gene on chromosome 22 and an ETS family gene, most frequently *FLI1* on chromosome 11, generating the fusion oncogene *EWS-FLI1*.^8–12^ This fusion protein acts as an aberrant transcription factor, driving the expression of genes that promote oncogenesis and tumor progression. One critical target of EWS-FLI1 is the Schlafen family member 11 (*SLFN11*) gene.^13^ EWS-FLI1 binds at regulatory regions adjacent to the transcription start site of the *SLFN11* promoter, acting as a positive regulator and leading to elevated *SLFN11* expression in EWS cells.^13,14^

*SLFN11* enhances sensitivity to DNA-damaging agents (DDAs) by blocking replication in the presence of DNA damage.^15–19^ Commonly used agents in EWS treatment like irinotecan, temozolomide, and lurbinectedin exploit this vulnerability.^3,20^ Prior studies demonstrated that combining PARP inhibitors with irinotecan and temozolomide induced complete and sustained tumor regression in over 80% of murine models.^3^ Recent preclinical data further showed that co-inhibition of PARP and topoisomerase I enhances the therapeutic efficacy of radiation in EWS.^21^ However, intrinsic and acquired resistance remain significant challenges, especially in patients with low or absent *SLFN11* expression.^3,22^ While high *SLFN11* expression is a hallmark of most EWS cell lines,^14^ patient tumors exhibit heterogeneity, and *SLFN11* knockout (*SLFN11^−/−^*) has been implicated in DDA resistance across multiple cancer types.^22^ In preclinical EWS models, *SLFN11^−/−^* induces chemoresistance both *in vitro* and *in vivo*, highlighting its pivotal role in modulating therapeutic efficacy.^22^ While its function in potentiating DDA-induced cytotoxicity is well established, its contribution to treatment response beyond the DNA damage repair pathway remains poorly understood.

Cancer cells rewire their metabolism to sustain rapid growth, survive under stress, and resist therapies by generating key metabolic substrates.^23–25^ This metabolic reprogramming not only fuels tumor progression but also contributes to chemoresistance by supporting redox balance, DNA repair, and facilitating drug inactivation or efflux.^26,27^ In this study, we investigated the metabolic pathway alterations associated with chemoresistance in *SLFN11^−/−^* EWS cell lines and patient derived xenograft model. Using integrated transcriptomic, metabolomic profiling, we identified a metabolic shift in *SLFN11^−/−^* EWS characterized by enhanced glycerophospholipid (GPL) biosynthesis. This reprogramming is driven by the suppression of glycerol-3-phosphate dehydrogenase (*GPD2*), a key mitochondrial enzyme that oxidizes glycerol-3-phosphate (G3P) to support mitochondrial respiration. Further, we found that therapeutic inhibition of GPL biosynthesis enhances the efficacy of DDA (SN-38) in *SLFN11^−/−^* EWS cells, providing a combination strategy to overcome chemoresistance.

These findings highlight a metabolic vulnerability associated with *SLFN11* loss and suggest that targeting GPL biosynthesis could help overcome DDA resistance. This work lays the groundwork for broader studies exploring metabolism-based strategies to counter chemoresistance.

## Results

### *SLFN11* expression is upregulated in EWS

Cancer Cell Line Encyclopedia (CCLE) is a compilation of gene expression, chromosomal copy number and next-generation sequencing data from 947 human cancer cell lines.^14,28^ Transcriptomic analysis from the CCLE reveals that *SLFN11* is broadly expressed across multiple cancer types, with EWS showing consistently elevated expression levels (Figure 1a). The Cancer Dependency Map (DepMap) is a functional genomics resource that integrates CRISPR and RNAi screening data with molecular profiles across diverse cancer cell lines to identify essential genes and lineage-specific dependencies.^29^ Analysis of EWS data derived from DepMap functional genomics further revealed a higher gene effect score (Chronos score) of *SLFN11* (*p*FDR = 1.4e-12) (Figure 1b). The high Chronos score suggests that *SLFN11* knockout does not compromise cell viability. Previous analyses of EWS patient cohorts and comparative datasets from The Cancer Genome Atlas Program (TCGA) have demonstrated that higher *SLFN11* expression is associated with improved therapeutic sensitivity and patient survival.^13^ Our analysis of the EWS cohort from the Ewing Sarcoma Cell Line Atlas (ESCLA)^30^ database also revealed that elevated *SLFN11* expression is associated with improved overall (*p* = 1.23e-3) and event-free (*p* = 6.85e-4) patient survival (Figure 1c-d). The ESCLA is a multi-omics resource of 18 EWS cell lines with inducible *EWSR1-ETS* knockdown, enabling analysis of fusion-driven gene regulation and molecular heterogeneity.^30^ Together, these analyses reveal that *SLFN11* is highly expressed in EWS and higher expression levels are associated with improved therapeutic response and patient survival, while its loss does not impact cell viability.

**Figure 1.**
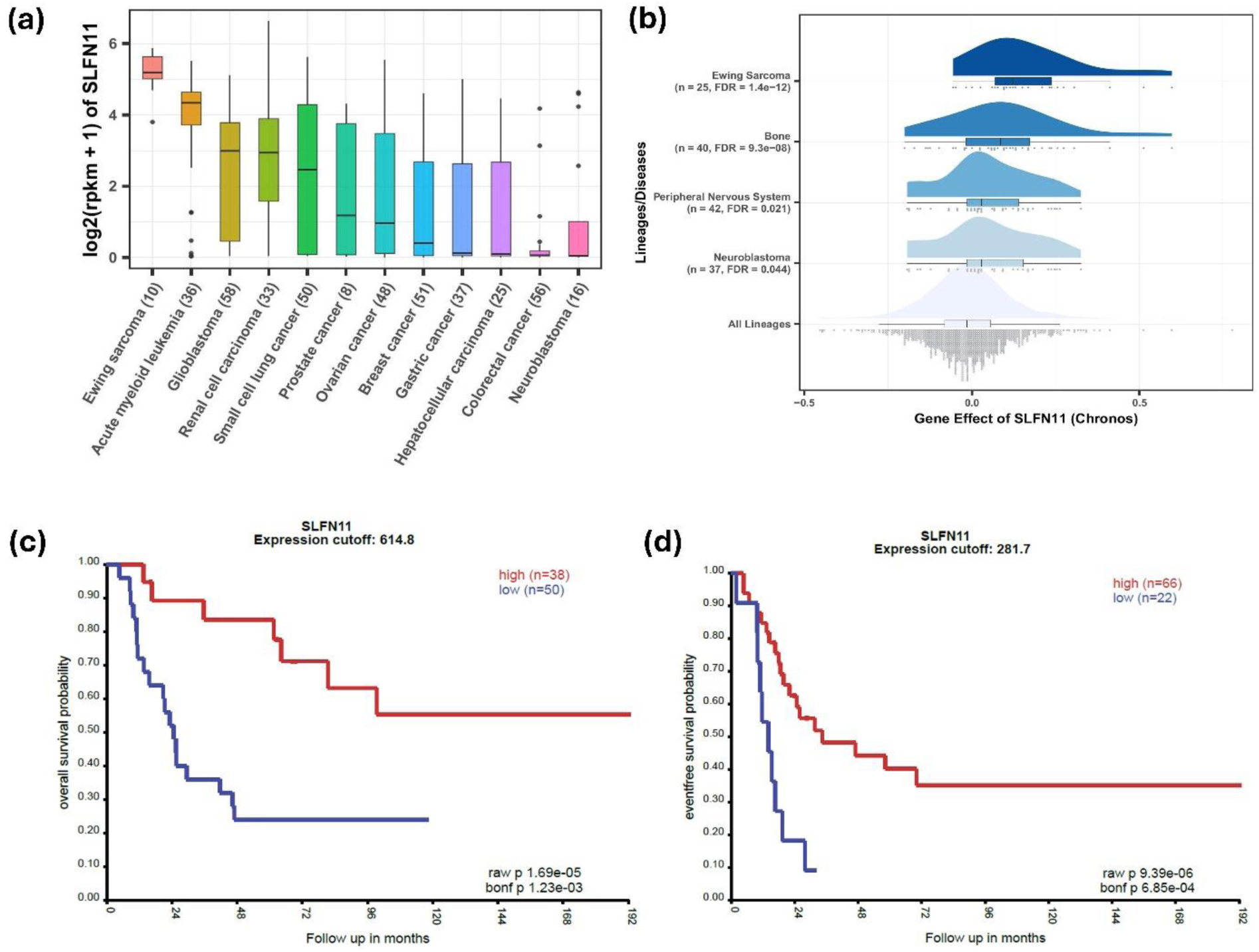
*SLFN11* is highly expressed in EWS and is associated with improved prognosis. **(a)** *SLFN11* expression across 428 primary tumors. Box-and-whisker plots show the distribution of mRNA expression for each subtype, ordered by the median *SLFN11* expression level (line), the inter-quartile range (box) and up to 1.5x the inter-quartile range (bars). Sample numbers (n) are indicated in parentheses. **(b)** Gene dependency analysis of *SLFN11* across tissue lineages using Chronos dependency scores from the DepMap project. **(c–d)** Kaplan–Meier survival analysis from the ESCLA database comparing patients with high (red line) versus low (blue line) *SLFN11* expression. Expression levels were dichotomized based on the median mRNA expression values. **(c)** Overall survival analysis of patients with high (n = 38) and low (n = 50) *SLFN11* expression using a median cutoff of 614.8. **(d)** Event-free survival analysis of patients with high (n = 66) and low (n = 22) *SLFN11* expression using a median cutoff of 281.7. n is number of patients.

### *SLFN11* knockout is associated with downregulation of the mitochondrial *GPD2*

To investigate *SLFN11*^−/−^ associated transcriptional changes more broadly, we performed RNA-sequencing (RNA-seq) analysis across patient derived EWS cell lines (Table S1). Principal component analysis (PCA) revealed clear segregation between WT and *SLFN11*^−/−^ cell lines along first principal component (PC1) (Figure S1a-e). RNAseq analysis further revealed a consistent downregulation of *GPD2* in the following EWS *SLFN11*^−/−^ cell lines: ES-8 (Log_2_FC= −6.5, *p*FDR = 5.45e-9) (Figure 2a), SK ES-1 (Log_2_FC= −1.33, *p*FDR = 4.5e-6) (Figure S1f), EW-8 (Log_2_FC= −0.73, *p*FDR = 5.27e−2) (Figure S1g), and RD ES-1 (Log_2_FC= −0.89, *p*FDR = 2.17e-7) (Figure S1h). However, *GPD2* expression was unaffected in CADO ES-1 cell line between WT and *SLFN11*^−/−^ cell lines (Figure S1i). The cytosolic isoforms of glycerol-3-phosphate dehydrogenase, either one of the *GPD1L* and *GPD1*, were differentially upregulated across the *SLFN11^−/−^* EWS cell lines; *GPD1L* expression was significantly elevated in ES-8 *SLFN11^−/−^* cell line (Log_2_FC = 0.65, *p*FDR = 2.14e-6) (Figure 2a), while *GPD1* expression was elevated in RD ES-1 *SLFN11^−/−^* cell line (Log_2_FC = 0.66, *p*FDR = 4.2e-2); (Figure S1h).

**Figure 2.**
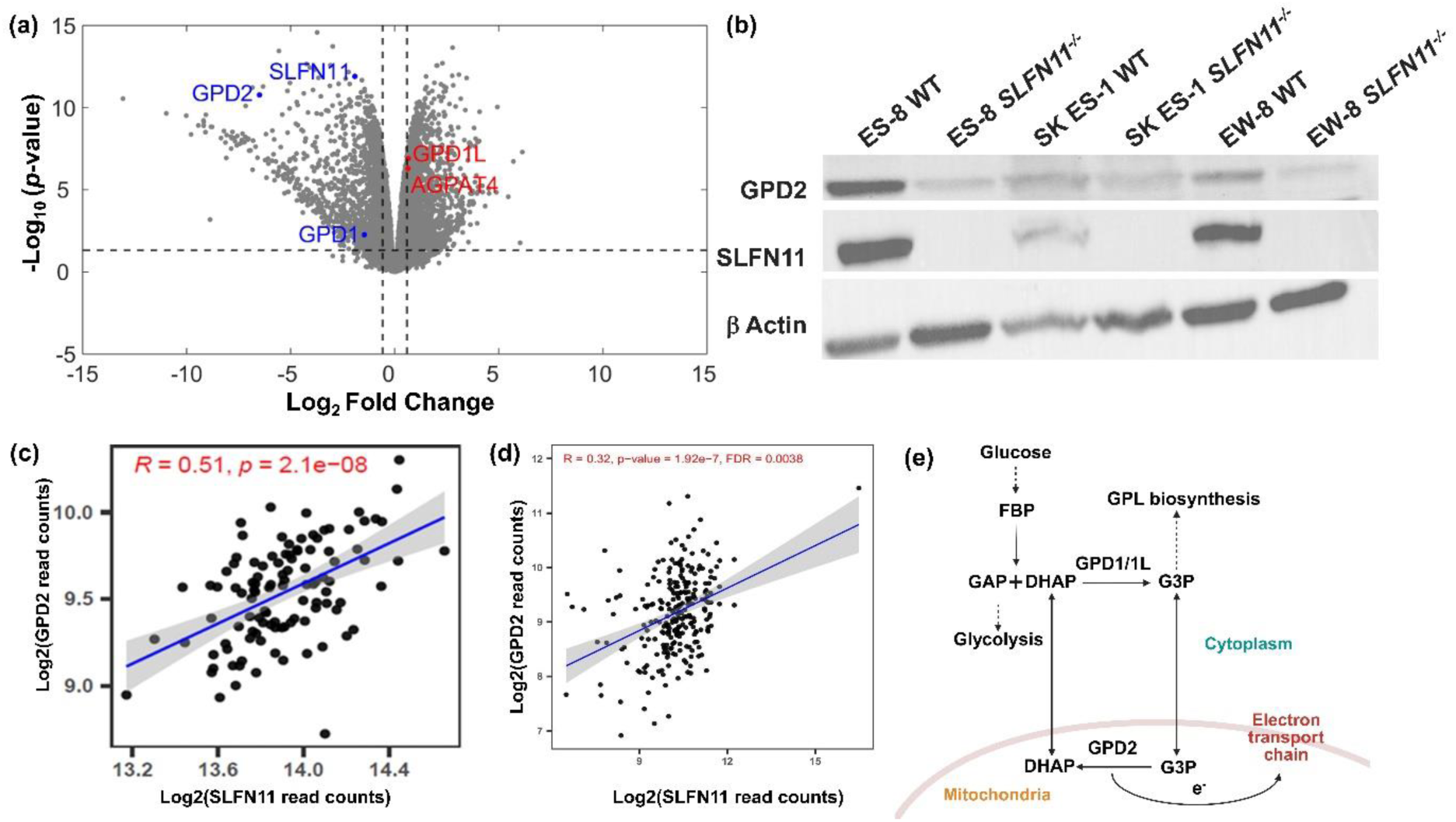
Knockout of *SLFN11* reprograms the G3PS through downregulation of mitochondrial *GPD2*. **(a)** Volcano plot showing differentially expressed genes in ES-8 *SLFN11*^−/−^ versus ES-8 WT cell line from RNA-seq analysis. Red dots indicate significantly upregulated genes involved in G3PS, while blue dots represent significantly downregulated genes. **(b)** Western blot analysis of GPD2 protein expression in ES-8 WT and *SLFN11*^−/−^, SK ES-1 WT and *SLFN11*^−/−^, and EW-8 WT and *SLFN11*^−/−^ cell lines. β-Actin serves as a loading control. **(c–d)** Correlation analysis between log₂-transformed *SLFN11* and *GPD2* read counts across EWS tumors in the **(c)** ESCLA dataset (R = 0.51, *p* = 1e-08) and **(d)** DepMap dataset (R = 0.32, *p* = 1.92e-07). R represents the Pearson correlation coefficient. Statistical significance was defined as p < 0.05. **(e)** Schematic of the G3PS showing cytosolic conversion of DHAP to G3P via *GPD1/1L*, followed by mitochondrial oxidation of G3P by *GPD2*, transferring electrons to the ETC to support OXPHOS. G3P also serves as the backbone for GPL biosynthesis in the cytosol. G3PS, glycerol-3-phosphate shuttle; DHAP, dihydroxyacetone phosphate; G3P, glycerol-3-phosphate; *GPD1/1L*, cytosolic glycerol-3-phosphate dehydrogenase; *GPD2*, mitochondrial glycerol-3-phosphate dehydrogenase; ETC, electron transport chain; OXPHOS, oxidative phosphorylation; GPL, glycerophospholipid.

To confirm the transcriptomics findings, we performed Western blot analysis of *GPD2* expression in selected EWS cell lines. As expected, *SLFN11*^−/−^ variants of ES-8, SK ES-1, and EW-8 exhibited a marked reduction in GPD2 protein, corroborating the transcriptomic findings (Figure 2b). Next, we examined publicly available datasets to assess broader expression correlations. Analysis of the ESCLA revealed a strong positive correlation between *SLFN11* and *GPD2* expression (Pearson R = 0.51, *p =* 2.1e-8) (Figure 2c). Similarly, across EWS cell lines in the DepMap database, *SLFN11* and *GPD2* expressions were positively correlated (Pearson R = 0.32, *p =* 1.92e-7) (Figure 2d).

These findings support a potential regulatory or functional relationship between *SLFN11* and *GPD2. GPD2* plays a central role in the glycerol-3-phosphate shuttle (G3PS), a key system connecting glycolysis, lipid metabolism, and redox homeostasis (Figure 2e).^31,32^ GPD2 catalyzes the oxidation of G3P to dihydroxyacetone phosphate (DHAP), a glycolytic intermediate. This interconversion transfers electrons to the electron transport chain (ETC). In the cytosol, *GPD1/GPD1L* reduces DHAP to G3P (Figure 2e).^31–33^ The G3PS enables redox coupling between cytosol and mitochondria by exchanging DHAP and G3P.

### *GPD2* downregulation elevates G3P levels in *SLFN11*-knockout cells

Building on these findings, we next investigated the metabolic consequences of *SLFN11* loss and the associated downregulation of *GPD2* by performing metabolomic profiling of *SLFN11*^−/−^ and WT EWS cell lines. To achieve this, we used LC/MS to profile a wide range of intracellular metabolites without predefined targets, enabling the identification of global metabolic alterations and pathway-level changes associated with *SLFN11* loss. Figure 3a illustrates the workflow for metabolic profiling. To maintain consistency and comparability across all samples, we performed batch effect removal and applied quantile normalization (Figure S2a-d). PCA was utilized to reduce data dimensionality and visualize patterns within the metabolomic profiles (Figure S2e-f). The PCA results revealed a clear separation between the *SLFN11*^−/−^ and the WT cell lines, indicating distinct metabolic states induced by *SLFN11* knockout in ES-8 (Figure S2e) and SK ES-1 (Figure S2f) cell lines.

**Figure 3.**
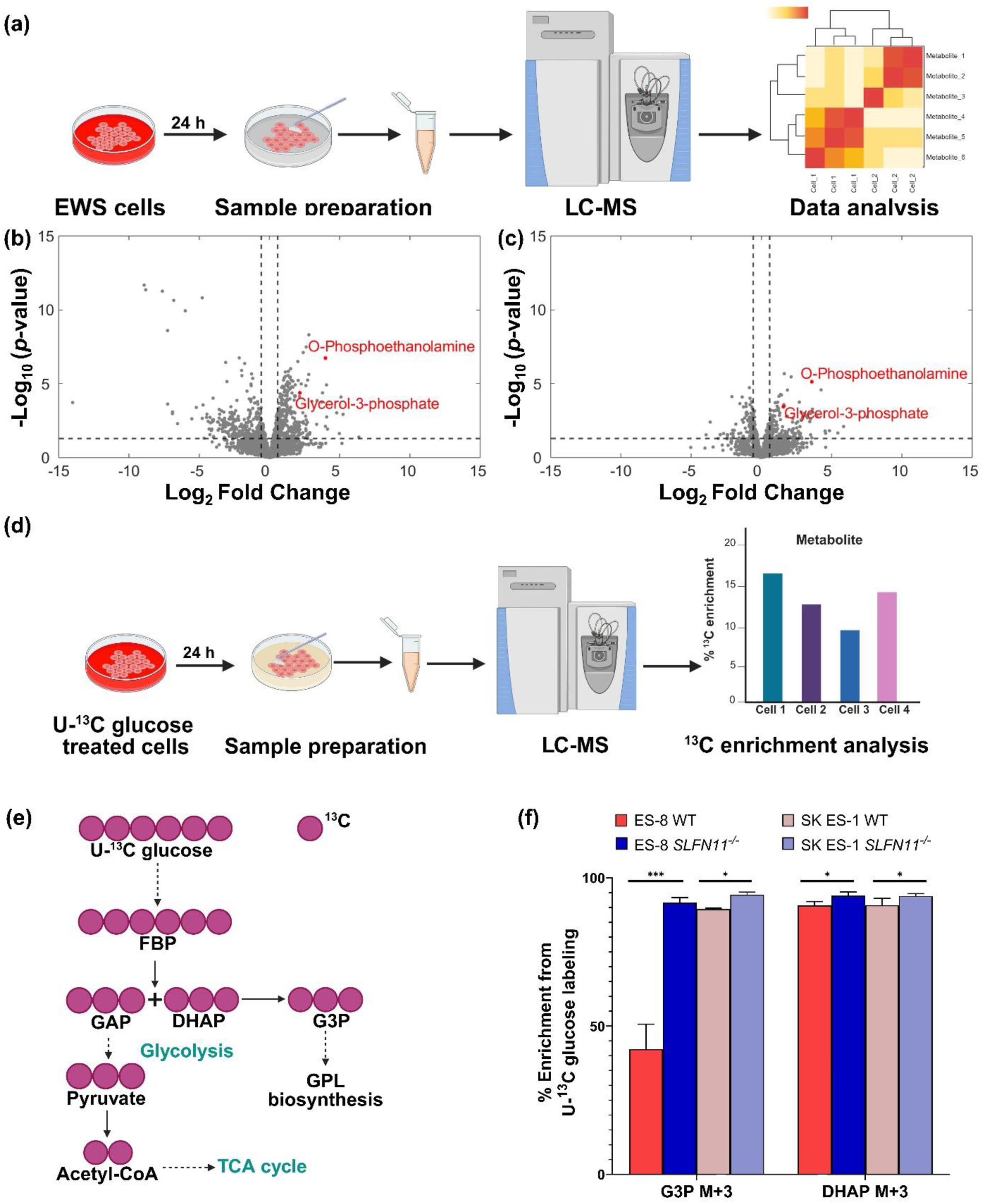
*SLFN11* knockout promotes G3P accumulation in EWS cells. **(a)** Schematic overview of the metabolomics workflow. Cells were harvested, extracted, and analyzed by LC-MS to identify metabolic alterations and differential metabolite abundance between cell lines. **(b-c)** Volcano plots illustrating differentially abundant metabolites between *SLFN11^−/−^* and WT cells in **(b)** ES-8 and **(c)** SK ES-1 cell lines. Red dots indicate significantly upregulated metabolites involved in GPL biosynthesis. **(d)** U¹³C glucose tracing to assess ¹³C enrichment in downstream metabolic intermediates. LC-MS analysis was performed to quantify isotopic labeling across cell lines. **(e)** Schematic showing U¹³C glucose-derived carbon incorporation into glycolysis and GPL biosynthesis. Purple circles represent ¹³C-labeled carbon atoms. **(f)** Bar graph showing percentage differences in ¹³C enrichment of the M+3 isotopologue for DHAP and G3P in WT and *SLFN11^−/−^* ES-8 and SK ES-1 cell lines following 24 h labeling with U¹³C glucose. G3P, glycerol-3-phosphate; DHAP, dihydroxyacetone phosphate. Data represent mean ± SD; n = 5. Statistical analysis was performed using paired two-tailed Student’s *t*-test for each comparison. *** *p* < 0.001, * *p* < 0.05. SD, standard deviation; n, number of replicates.

To further investigate the metabolic changes associated with *SLFN11* knockout, we generated volcano plots based on unsupervised clustering analysis (Figure 3b-c, Table S2-S3). These plots highlighted a considerable number of upregulated and downregulated metabolites in *SLFN11*^−/−^ cell line compared to WT cell line (Figure 3b-c, Table S2-S3). Notably, G3P showed a marked increase in both ES-8 (Log_2_FC=2.28, *p*FDR = 1.43e-6) (Figure 3b, Table S2) and SK ES-1 (Log_2_FC=1.5, *p*FDR = 7e-4) (Figure 3c, Table S3) *SLFN11*^−/−^ cell lines compared to their respective WT cell lines. To validate these findings, we performed stable isotope tracing using U-¹³C glucose. This approach enables the quantification of glycolytic flux into G3P, linking glucose metabolism to GPL synthesis as illustrated in Figure 3e. Increased ¹³C enrichment in G3P reflects enhanced channeling of glucose-derived carbon toward lipid biosynthesis. Uniformly labeled ¹³C-glucose generates predominantly M+3 isotopologues of G3P, signifying complete incorporation of three ¹³C atoms into its carbon backbone.^34^ Using this approach, we observed that ES-8 *SLFN11*^−/−^ exhibited a 2.17-fold higher ^13^C labeling in G3P M+3 isotopologue compared to ES-8 WT cell line (*p* = 5.7e-4), (Figure 3f, Table S4) confirming elevated G3P production. Similarly, SK ES-1 cell line exhibited an increase in G3P labeling (*p* = 1.52e-4) in the *SLFN11*^−/−^ cell line (Figure 3f, Table S5). DHAP was upregulated in ES-8 *SLFN11*−/−cell line by 1.04-fold (*p* = 4e-2) and in SK ES-1 *SLFN11^−/−^* cell line by 1.03-fold (*p* = 1.63e-2) compared to their respective WT cell lines (Figure 3f, Table S4-S5).

We further observed an upregulation of O-phosphoethanolamine in ES-8 *SLFN11^−/−^* (Log_2_FC=3.9, *p*FDR = 9.07e-10) (Figure 3b, Table S2) and SK ES-1 *SLFN11^−/−^* (Log_2_FC=3.63, *p*FDR = 1.15e-8) (Figure 3c, Table S3) cell lines compared to corresponding WT cell lines. O-Phosphoethanolamine is a key intermediate in phosphatidylethanolamine biosynthesis.^35^

Collectively, these findings reveal that GPD2 downregulation in *SLFN11* knockout cells leads to G3P accumulation and altered metabolic flux involving the G3PS. Previous studies have reported that GPD2 knockdown results in G3P accumulation and facilitates the synthesis of complex lipids.^32^ Our findings of elevated G3P levels suggest a potential shift toward lipid biosynthesis in *SLFN11^−/−^* EWS cells, as G3P serves as the backbone for the synthesis of key GPL species, including PE, phosphatidylcholine (PC), phosphatidylinositol (PI), phosphatidylserine, phosphatidylglycerol (PG), cardiolipin, phosphatidic acid (PA), diacylglycerol (DAG), and triacylglycerol via sequential acylation with fatty acids.^31,32,36^

### *SLFN11* knockout promotes GPL biosynthesis via *GPD2* downregulation

Based on the link between *SLFN11* loss, *GPD2* downregulation, and G3P accumulation, we next sought to identify the metabolic pathways affected by *SLFN11* knockout. To achieve this, we performed metabolite set enrichment analysis (MSEA) using MetaboAnalyst 6.0.^38,39^ The analysis identified GPL biosynthesis as a key pathway upregulated with *SLFN11* knockout in both ES-8 (Figure 4a, Table S6) and SK ES-1 (Figure S2g, Table S7) cell lines. While the findings in SK ES-1 achieved statistical significance (*p* = 9.30e-3), the results in ES-8 exhibited a trend.

**Figure 4.**
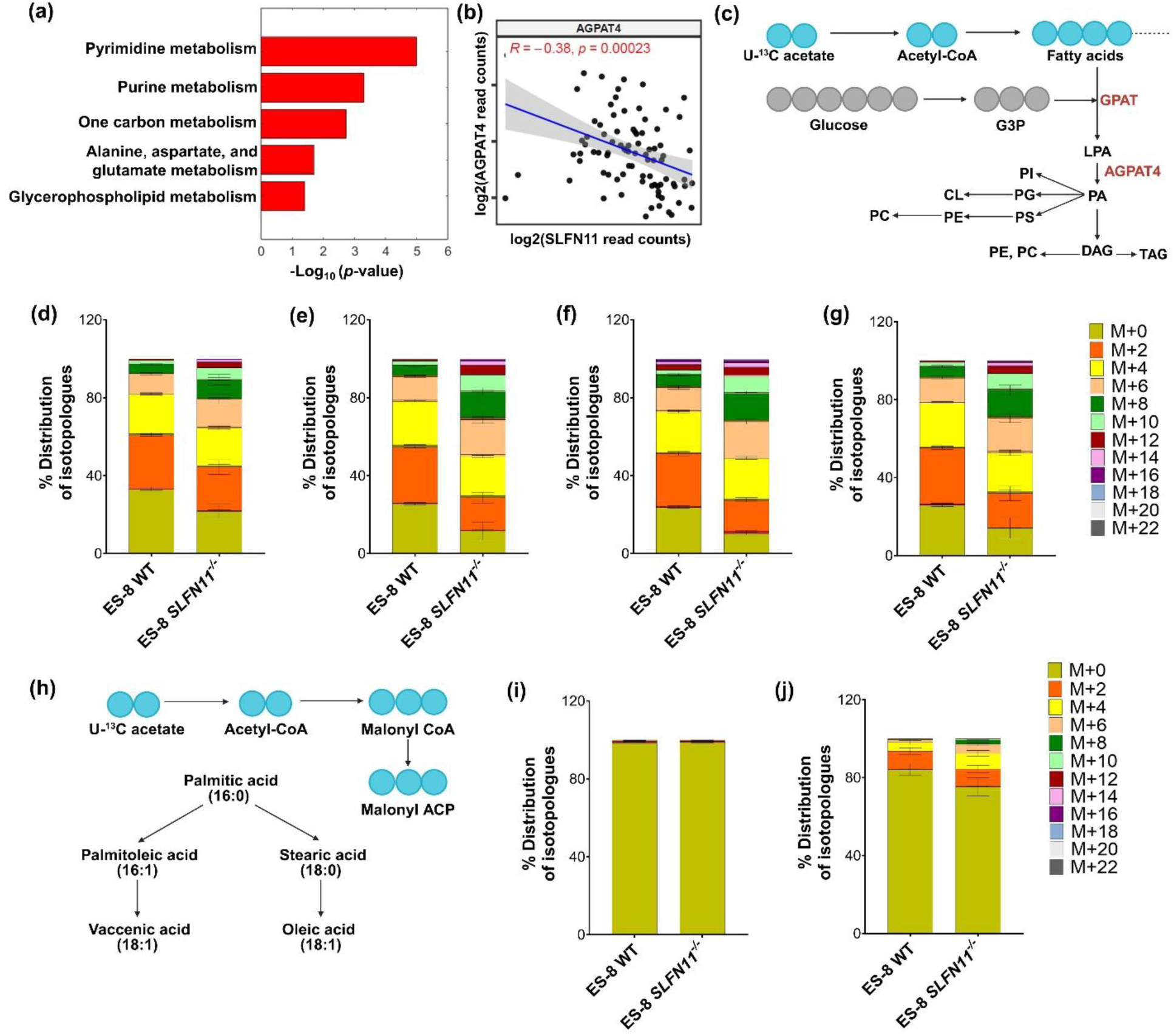
*SLFN11* knockout increases GPL biosynthesis in EWS. **(a)** MSEA-based pathway enrichment analysis identifying upregulated metabolic pathways in ES-8 *SLFN11^−/−^* cell line. Pathways with −log₁₀(FDR) ≥ 0.5 were considered enriched and ranked by significance. **(b)** Correlation analysis of log₂-transformed *SLFN11* and *AGPAT4* expression across EWS tumors using patient-derived microarray data^37^ from the Gene Expression Omnibus (GEO) database. (R = −0.38, *p* = 2.3e-04). R represents the Pearson correlation coefficient. Statistical significance was defined as *p* < 0.05. **(c)** Schematic illustrating U¹³C acetate incorporation into the GPL biosynthesis pathway. Cyan circles represent ¹³C-labeled acetate-derived carbons incorporated into fatty acid chains. Gray circles denote the synthesis of G3P backbone. Enzymes involved in GPL biosynthesis are highlighted in maroon. **(d-g)** Isotopologue distribution of U¹³C acetate-labeled GPL species: **(d)** PE (34:1), **(e)** PC (34:1), **(f)** PG (34:1), and **(g)** PI (34:1) in ES-8 WT and *SLFN11^−/−^* cells. Bars represent the relative abundance of individual mass isotopologues (M+2 to M+22), corresponding to successive incorporation of ¹³C-labeled acetate units during GPL synthesis. A color scale denoting isotopologue species (M+0 to M+22) is included alongside the figure. Data represents mean ± SD (n = 5). Statistical comparisons were performed using paired two-tailed Student’s *t*-test for each comparison. SD, standard deviation; n, number of replicates. **(h)** Schematic of U¹³C acetate tracing into *de novo* fatty acid synthesis. Cyan circles represent ¹³C-labeled acetate-derived carbons incorporated during elongation. The diagram illustrates labeling flow into saturated and monounsaturated fatty acid species. **(i-j)** Distribution of U¹³C acetate-labeled isotopologues in **(i)** palmitic acid and **(j)** oleic acid in ES-8 WT and *SLFN11^−/−^* cells. Color code for isotopologues (M+0 to M+22) is shown adjacent to panel. Bars represent mean ± SD (n = 5). Statistical comparisons were performed using paired two-tailed Student’s *t*-test for each comparison. SD, standard deviation; n, number of replicates.

Transcriptomics analysis demonstrated significant upregulation of *AGPAT4* (Log_2_FC = 0.63, *p*FDR = 6.37e-6), a critical gene involved in GPL biosynthesis,^32,40,41^ in the ES-8 *SLFN11*^−/−^ cell line (Figure 2a). To validate this observation in patient-derived data, we analyzed EWS tumor microarray profiles from the Gene Expression Omnibus (GEO) database, which revealed a significant inverse correlation between *SLFN11* and *AGPAT4* expression (Pearson R = −0.38, *p =* 2.3e-4) (Figure 4b).

To further investigate the observed upregulation of GPL biosynthesis, we performed stable isotope tracing using U-¹³C acetate. Acetate serves as a key carbon source for fatty acid biosynthesis by supplying two-carbon units through acetyl-CoA. U-¹³C acetate labeling method enables quantification of carbon incorporation into GPL.^42^ Each acetate unit provides two ¹³C atoms, producing M+2, M+4, M+6, and higher isotopologues through stepwise incorporation during fatty acid elongation as illustrated in Figure 4c. Elevated levels of these isotopologues indicate enhanced *de novo* lipid synthesis from acetate.^42^ Given the significantly higher ¹³C-glucose-derived G3P labeling observed in ES-8 cells (Figure 3f), we selected this cell line for further analysis. Patient-derived ES-8 cell lines were chosen for U-¹³C acetate tracing due to their consistent metabolic profile, making them an optimal system to assess GPL synthesis. LC-MS analysis revealed significant incorporation of ¹³C-labeled acetate-derived carbons into GPL species in ES-8 *SLFN11^−/−^* compared to ES-8 WT cell line (Figure 4d-g). The total ¹³C labeling of PE was higher in *SLFN11^−/−^* (78%) compared to WT ES-8 cell line (67%) (*p* = 2.72e-03) (Fig. 4d). Similarly, in PC, 88.4% labeling was observed in *SLFN11^−/−^* cell line, compared to 74.4% in WT (*p =* 3.18e-3) (Figure 4e). In PG ∼90% of the lipid pool was labeled in *SLFN11^−/−^* cell line versus 76% in WT (*p =* 2.21e-9) (Figure 4f). PI showed 86% labeling in *SLFN11*^−/−^ versus 74% in WT (*p =* 1.03e-2) (Figure 4g). These findings suggest that *SLFN11* knockout enhances ^13^C-acetate flux into membrane lipid synthesis, indicating a shift toward increased reliance on acetate for *de novo* GPL biosynthesis. The distribution of all isotopologues of these lipid species is presented in Figure 4d-g and Table S8.

Additionally, we assessed fatty acid labeling using U¹³C acetate, as illustrated in Figure 4h. This tracer enables evaluation of fatty acid elongation by tracking the incorporation of ¹³C-labeled acetate units. *SLFN11^−/−^* cells exhibited increased labeling in monounsaturated fatty acids (MUFAs). No significant difference of ¹³C labeling in palmitate (saturated fatty acid) was observed between ES-8 *SLFN11^−/−^* and ES-8 WT cell lines (Figure 4i), whereas increased labeling of oleate (unsaturated fatty acid) was seen in ES-8 *SLFN11^−/−^* cell line (∼25%) compared to ES-8 WT cell line (∼15%) (Figure 4j) (*p* = 1.0e-2).

Taken together, these results highlight the metabolic reprogramming driven by *SLFN11* knockout, with a marked shift toward unsaturated lipid biosynthesis pathways.

### *SLFN11* knockout sensitizes EWS cells to combined DNA damage and lipid biosynthesis inhibition

To explore whether the elevated GPL pathway in *SLFN11^−/−^* cells could be therapeutically exploited, we performed a Bivariate Response to Additive Interacting Doses (BRAID)-format drug combination screen.^43,44^ This screen was conducted in ES-8 WT and *SLFN11^−/−^* cells to evaluate the efficacy of combining DNA-damaging agents with lipid biosynthesis inhibitors.^44,45^ BRAID fits an eight-parameter nonlinear model to describe the combined dose response behavior.^43,44^ SN-38, a potent topoisomerase I inhibitor^46^ with *SLFN11*-dependent activity (Figure 5a),^16,22^ was used as the anchor drug and combined with FSG67, a selective GPAT1 inhibitor. GPAT1 is a key enzyme in *de novo* GPL synthesis that supports membrane lipid production (Figure 5b).^32^

**Figure 5.**
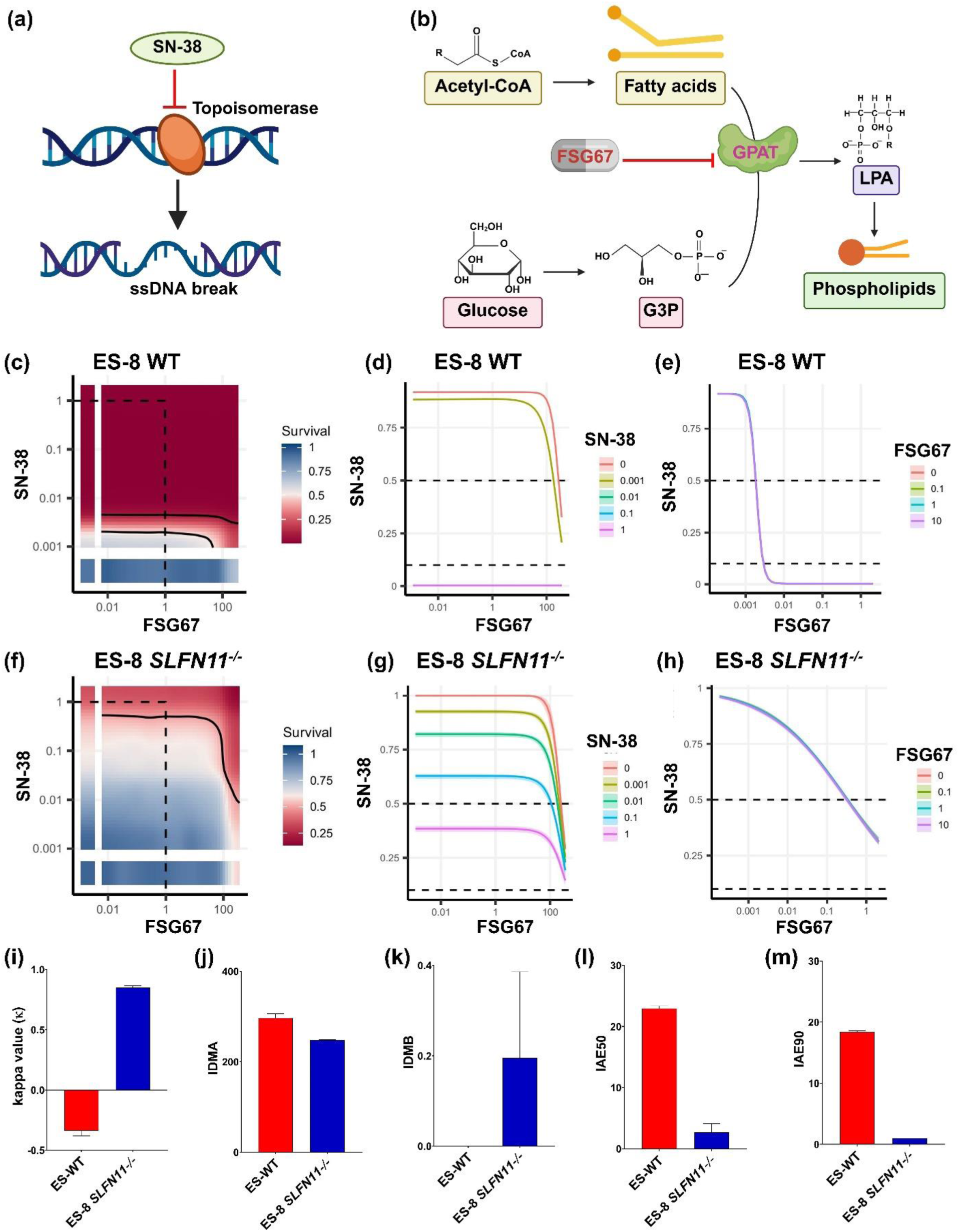
*SLFN11* knockout sensitizes EWS cells to combined DNA damage and *GPAT1* inhibition. **(a)** Schematic representation of SN-38 mechanism of action. SN-38 inhibits topoisomerase I, stabilizing DNA single-strand breaks (ssDNA break) during replication and inducing cytotoxicity, particularly in *SLFN11*-expressing cells. **(b)** Schematic of FSG67 mechanism of action. G3P, derived from glucose, is converted to phosphatidic acid (PA) by glycerol-3-phosphate acyltransferase (*GPAT*). FSG67 inhibits *GPAT1*, impairing *de novo* GPL biosynthesis and limiting membrane lipid production **(c– m)** BRAID analysis of ES-8 WT and *SLFN11^−/−^* cells following 72 h *in vitro* co-treatment with SN-38 and FSG67. **(c)** Representative response surface plot showing survival probability across concentration combinations in ES-8 WT cell line. The red and blue color scales indicate low and high proliferation, respectively, relative to DMSO-treated cells. The dotted line encompasses the region in the response surface where each drug is ≤1 μmol/L and shown for reference. **(d)** Representative SN-38 potentiation of FSG67 dose– response in ES-8 WT cell line. **(e)** Representative FSG67 potentiation of SN-38 dose– response in ES-8 WT cell line. **(f)** Representative response surface plot showing survival probability across concentration combinations in *SLFN11^−/−^* cell line. **(g)** Representative SN-38 potentiation of FSG67 dose–response in *SLFN11^−/−^* cell line. **(h)** Representative FSG67 potentiation of SN-38 dose–response in *SLFN11^−/−^* cell line. **(i–m)** Combination treatment metrics in ES-8 WT and *SLFN11^−/−^* cells (n = 2 biological replicates, each with 3 technical replicates): **(i)** Kappa interaction coefficient (κ) for SN-38 and FSG67 (κ ≈ 0: additive; κ > 0: synergism; κ < 0: antagonism). **(j)** IDMA: EC_50_ of the anchor drug SN-38. **(k)** IDMB: EC_50_ of the partner drug FSG67. **(l)** IAE50: combination dose required for 50% cell death. **(m)** IAE90: combination dose required for 90% cell death.

We assessed cell survival following 72 h of SN-38 and FSG67 co-treatment using CellTiter-Glo followed by BRAID modeling (Figure 5c-m). Synergy between SN-38 and FSG67 was evaluated using the κ (kappa) parameter, where κ > 0 indicates synergy, κ = 0 represents additivity, and κ < 0 indicates antagonism.^43,44^ Combination efficacy was further quantified using IAE50 and IAE90 (Index of Achievable Efficacy), which represent the proportion of tested concentration pairs achieving at least 50% or 90% inhibition, respectively, based on the fitted BRAID response surface.^43,44^

In ES-8 WT cells, SN-38 was highly effective as a single agent (Figure 5e,k; Table S9), consistent with prior observations of *SLFN11*-dependent cytotoxicity.²⁴ However, co-treatment with FSG67 in these cells resulted in limited additional benefit, as indicated by a predominantly high-survival region in the BRAID surface and negative κ values (−0.31 and −0.37) with narrow confidence intervals excluding zero, indicative of statistically significant antagonism (Figure 5c,i; Table S9). Dose response plot demonstrated that FSG67 alone showed minimal cytotoxicity across the tested range (Figure 5d,j; Table S9) and even in combination with SN-38, its contribution to overall efficacy in WT cells remained minimal.

In contrast, *SLFN11^−/−^* cells displayed marked synergy between SN-38 and FSG67. BRAID modeling revealed a broad region of low survival probability upon co-treatment, and κ values were consistently positive (0.86 and 0.84) (Figure 5f,i; Table S9), with confidence intervals excluding zero (Table S9), confirming strong synergistic interaction (Figure 5f,i; Table S9). Dose-response plots showed pronounced potentiation of each drug by the other, particularly at higher concentrations, resulting in a dose-dependent reduction in cell viability (Figure 5g-h). Despite this synergy, overall combination efficacy in *SLFN11^−/−^* cells was modest, with IAE50 values of 1.72 and 3.70, and IAE90 values approaching 1.00 (Figure 5l-m; Table S9). WT cells exhibited much higher absolute efficacy driven by SN-38 alone, with IAE50 and IAE90 values, approximately 23 and 18, respectively.

FSG67 EC50 values (IDMB) were slightly lower in *SLFN11^−/−^* cells (246.47 and 248.33 μM) compared to WT (289.99 and 303.50 μM), indicating a mild increase in sensitivity to lipid biosynthesis inhibition (Figure 5j, Table S9). As the maximal inhibitory effect was not reached within the tested concentration range, the EC50 values likely underestimate the true potency of FSG67. SN-38 exhibited reduced potency (IDMA) in ES-8 *SLFN11^−/−^* cells compared to the WT (Figure 5k) consistent with previous observation,^22^ demonstrating *SLFN11*-dependent sensitivity to DNA-damaging agents.

Collectively, these data support the conclusion that *SLFN11* knockout selectively sensitizes ES-8 cells to combined DNA damage and lipid biosynthesis blockade, revealing a therapeutically targetable metabolic vulnerability.

### *SLFN11* loss alters choline metabolism in EWS tumor xenografts

To determine whether the lipid metabolic reprogramming observed in vitro is reflected in vivo, we analyzed xenografted tumors derived from ES-8 WT and ES-8 *SLFN11^−/−^* cells. NMR spectroscopy was performed on metabolites extracted from these tumors in athymic nude mice, as outlined in Figure 6a. ^47–49^ NMR spectroscopy enables quantification of water-soluble metabolites, allowing evaluation of GPL metabolic alterations in tumor extracts.^48^ Figure 6b and Figure 6c show representative ¹H NMR spectra of WT and *SLFN11^−/−^* cells, respectively. Data averaged from three tumors per group revealed a consistent increase in phosphocholine (PCh) (*p* = 0.18) and total choline (Choline) (*p* = 0.04) levels in the ES-8 *SLFN11^−/−^* group compared to ES-8 WT tumors (Figure 6d). In contrast, Glycerophosphocholine (GPC) levels exhibited a downward trend (*p* = 0.39) in ES-8 *SLFN11^−/−^* tumors (Figure 6d). However, none of these changes reached statistical significance. Notably, the PCh/GPC ratio was significantly elevated in the *SLFN11^−/−^* group (*p* = 0.03) (Figure 6e). In the Kennedy pathway, PCh contributes to GPL synthesis, while GPC results from GPL breakdown, making the PCh/GPC ratio indicative of GPL turnover (Figure 6f). This shift indicates enhanced choline-driven GPL biosynthesis in *SLFN11^−/−^* xenografts.

**Figure 6.**
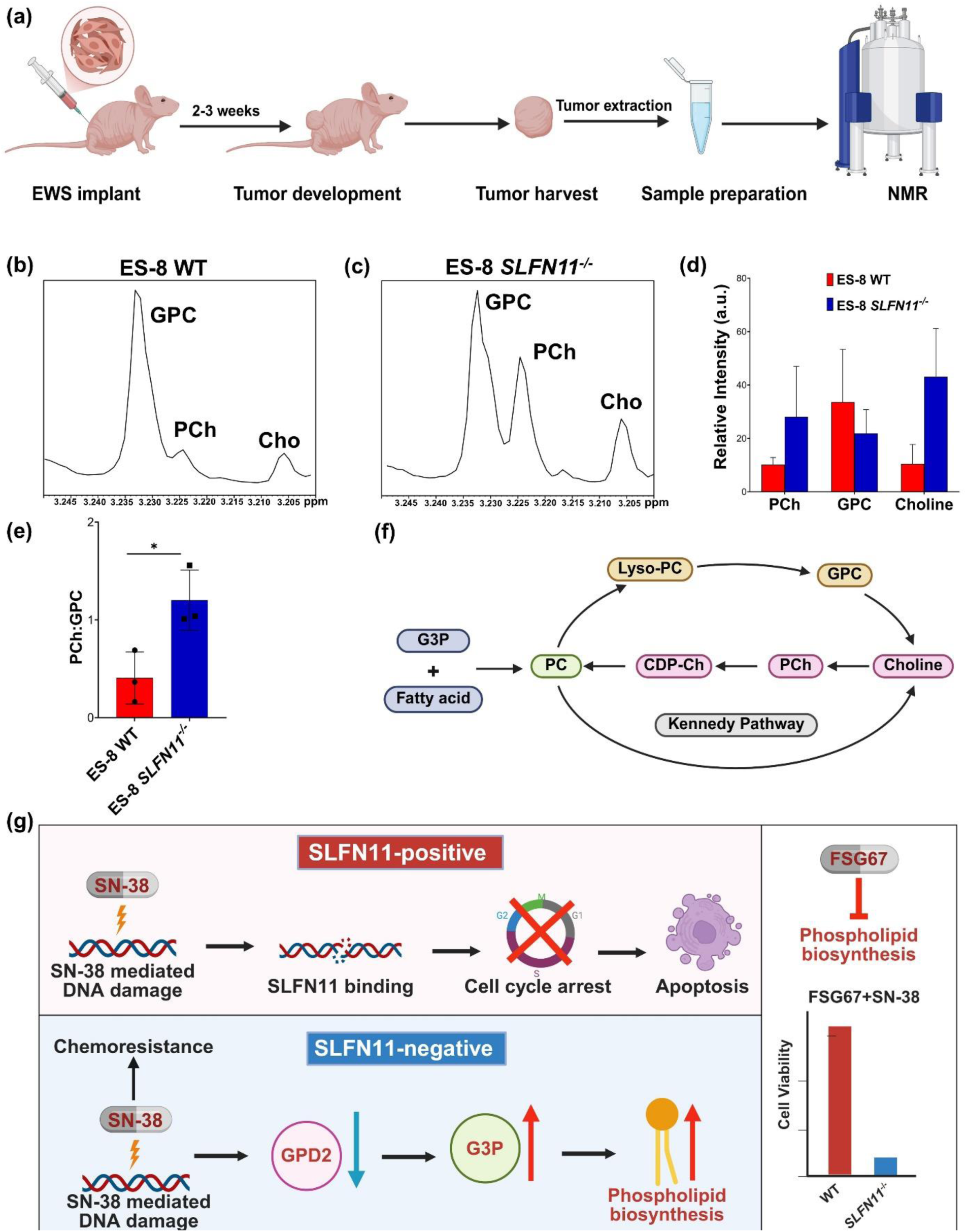
*Ex vivo* ¹H NMR profiling reveals altered choline metabolism in *SLFN11^−/−^* EWS tumors. **(a)** Schematic of *ex vivo* ¹H NMR (600 MHz) analysis in athymic nude mice. EWS cells were implanted and excised tumors were extracted for metabolic profiling. **(b-c)** Representative ¹H NMR spectra of choline metabolite obtained from water-soluble tumor extract from **(b)** ES-8 WT and **(c)** *SLFN11^−/−^* xenografts. Spectra are expanded to display signals from 3.20 to 3.25 ppm. Metabolites involved in choline metabolism including GPC (3.235 ppm), (PCh, 3.226 ppm), and free choline (3.208 ppm), highlighted with arrows. **(d)** Bar graph showing quantification of GPC, PCh, and total choline levels in WT and *SLFN11^−/−^* ES-8 tumor xenografts. Data represent mean ± SD (n = 3). Statistical significance was determined using paired two-tailed *t*-test. **(e)** Quantification of PCh/GPC ratio in ES-8 WT and *SLFN11^−/−^* xenograft tumors. Data represent mean ± SD (n = 3). Statistical significance determined using paired two-tailed *t*-test (*p < 0.05). SD, standard deviation; n, number of replicates. **(f)** Schematic illustrating the Kennedy pathway (pink) involved in choline metabolism, where choline is converted to PC via PCh and CDP-Ch. An alternate route involving G3P and fatty acids (blue) also contributes to PC generation. GPC and Lyso-PC degradation branches are shown in beige. PC, phosphatidylcholine; CDP-Ch, CDP-choline; PCh, phosphocholine; GPC, glycerophosphocholine; G3P, glycerol-3-phosphate. **(g)** Schematic summary of *SLFN11*-loss-induced metabolic rewiring in EWS.

Together, our findings show that *SLFN11* loss causes metabolic changes in EWS, including *GPD2* downregulation, G3P accumulation, and a higher PCh:GPC ratio. These changes suggest increased GPL biosynthesis in *SLFN11*^−/−^ cells. This creates a metabolic vulnerability that can be targeted by combining DNA-damaging agents with lipid biosynthesis inhibitors. The proposed mechanism of metabolic reprogramming due to *SLFN11* loss is depicted schematically in Figure 6g.

## Discussion

*SLFN11* is a critical modulator of cancer cell sensitivity to DDAs.^13,16–18,50,51^ In response to DNA damage, *SLFN11* is recruited to replication forks via RPA1^16–18^ and interacts with the MCM3^16,17^. This interaction promotes chromatin opening and induces irreversible replication arrest without disrupting the loading of CDC45^16,17^ or PCNA.^17^ This mechanism contrasts with ATR-mediated fork slowing and allows *SLFN11* to exert a dominant cytotoxic effect under genotoxic stress.^52,53^ Notably, *SLFN11* functions by degrading CDT1 through DDB1–CUL4CDT2 E3 ubiquitin ligase complex, thereby blocking replication reactivation after damage.^18^ Expression of *SLFN11* strongly correlates with responsiveness to a broad spectrum of DDAs, including topoisomerase I/II inhibitors, alkylating agents, DNA synthesis inhibitors, and PARP inhibitors, across multiple cancer types.^3,16^ Despite its therapeutic relevance, *SLFN11* is frequently inactivated, most commonly by promoter hypermethylation or histone modifications.^16–18,51^ This loss leads to chemoresistance in approximately 50% of cancer cell lines.^18,51^ Elevated *SLFN11* expression is associated with improved tumor-free survival in EWS, underscoring its clinical relevance.^13^ Consistently, in our analysis of ESCLA, high *SLFN11* levels were strongly associated with improved overall and event-free survival. These findings support an association between *SLFN11* expression and improved clinical outcome and drug sensitivity in EWS. Additionally, our evaluation of DepMap functional dependencies and transcriptomic profiles across EWS cell lines further reinforced this association, highlighting the selective advantage of cells with *SLFN11* expression under DDA therapy. Although *SLFN11* is expressed in the majority of EWS tumors, a subset (∼10%) lacks detectable expression,^22,54^ which may contribute to reduced sensitivity to DDAs. This underscores the importance of identifying alternative therapeutic strategies or predictive markers for *SLFN11*-negative EWS.

Metabolic reprogramming is increasingly recognized as a hallmark of cancer and a key contributor to therapeutic resistance.^24^ Prior studies have leveraged metabolic characterization to uncover dependencies in oncogenic signaling and drug resistance across cancer types.^41,42,55–61^ Garrett et al. employed comprehensive metabolic profiling to characterize distinct metabolic programs across medulloblastoma subgroups, including alterations in purine and amino acid metabolism.^60^ Stable isotope tracing has further clarified nutrient utilization in both *in vitro* and *in vivo* models.^59^ DeBerardinis and colleagues demonstrated that mitochondrial respiration sustains tricarboxylic acid cycle activity and supports anabolic growth in non-small cell lung cancer (NSCLC), establishing mitochondrial glucose oxidation as a key feature of tumor metabolism.^41,59^ Additional studies from this group expanded on these insights by identifying a metabolic switch from *de novo* purine biosynthesis to salvage pathways under electron transport chain inhibition, driven by redox imbalance and augmented by purine nucleobase uptake in NSCLC.^58^ In a parallel study, the same group showed that glutamine fuels TCA cycle intermediates via both oxidative and reductive pathways in ccRCC models.^61^ Formate overflow from serine catabolism has been shown to promote tumor invasiveness.^55–57^ Acetate utilization, in turn, supports lipid synthesis as a compensatory adaptation to metabolic stress.^42^

Building on these insights, our integrated transcriptomic and metabolomic profiling of *SLFN11^−/−^* EWS models revealed a critical link between *SLFN11* loss–mediated chemoresistance and metabolic reprogramming, uncovering distinct vulnerabilities that may be exploited to overcome therapeutic resistance. A consistent feature was the downregulation of mitochondrial *GPD2*, a key component of the ETC and oxidative phosphorylation, ^32^ leading to impaired mitochondrial function and cytosolic accumulation of G3P, a precursor for *de novo* GPL biosynthesis.^36^ Previous studies have shown that *GPD2* downregulation leads to upregulation of cytosolic *GPD1*, which enhances G3P production.^32^ In glioblastoma, *GPD1* has been implicated in chemoresistance through its role in regulating GPL metabolism. Knockout of *GPD1* sensitizes glioblastoma cells to temozolomide, highlighting its contribution to therapy resistance.^62^ Our study showed that elevated G3P levels drive the synthesis of key GPL, including PE, PC, PG, and PI, alongside increased expression of *AGPAT4*, a critical gene in GPL biosynthesis.^52,53^ Multiple studies have shown that GPL synthesis rates are elevated during oncogenesis and tumor progression,^63–67^ including in lung,^65^ breast,^64,66^ colorectal,^67^ bladder, and renal cancers^68^ compared to normal tissue. Lesko *et al.* showed that lung cancer exhibits elevated *de novo* synthesis and turnover of phosphatidylethanolamine, indicating enhanced GPL metabolism in tumors.^65^ Increased GPL biosynthesis also generates signaling lipids such as DAG and PA, which activate pro-survival pathways like mTOR.^69^ *SLFN11* has been reported to suppress mTOR-driven tumorigenesis, suggesting that its loss may facilitate activation of lipid-sensitive survival signaling.^70^ In addition to their structural role, GPL regulates oncogenic signaling, redox homeostasis, ferroptosis susceptibility, and therapeutic resistance.^63,71^ Increased GPL biosynthesis supports membrane remodeling,^72^ promotes proliferation,^73^ and limits lipid peroxidation,^74^ helping tumor cells withstand metabolic and therapeutic stress.^63,71^ Our observation of enhanced GPL biosynthesis in *SLFN11^−/−^* EWS cells points to a potential role for lipid remodeling in sustaining chemoresistance within this population. Consistent with this metabolic shift, our *in vivo* metabolic analysis revealed an elevated PCh/GPC ratio in *SLFN11^−/−^* EWS tumors. A high PCh to GPC ratio reflects elevated PC biosynthesis through Kennedy pathway and has been associated with increased tumor aggressiveness, cell proliferation, and therapeutic resistance across multiple cancers.^48,49,75,76^ The elevated PCh/GPC ratio may serve as a functional biomarker of chemoresistance in EWS. Targeting *de novo* lipogenesis has been shown to overcome chemoresistance in several cancers.^77^ Inhibitors of key lipogenic enzymes such as FASN, ACC, and SCD1 enhance drug sensitivity in models of pancreatic, ovarian, prostate, breast, and liver cancer, highlighting lipid biosynthesis as a promising therapeutic target.^77^ Pharmacologic inhibition of GPAT1, a rate-limiting enzyme in GPL synthesis with FSG67 has shown preclinical efficacy by limiting GPL synthesis and suppressing tumor growth.^32^

Loss of *SLFN11* expression in EWS reduces the effectiveness of DDAs, limiting therapeutic response and contributing to treatment resistance.^13^ Previous studies have shown that *SLFN11*-expressing tumors respond robustly to SN-38 and other DDAs, whereas *SLFN11^−/−^* EWS shows resistance sensitivity, presenting a major therapeutic challenge.^3,22^ Our findings suggest that *SLFN11* loss not only contributes to chemoresistance but also drives metabolic adaptations that support survival under genotoxic stress. We hypothesized that dual targeting of DNA damage and lipid biosynthesis pathways could counteract chemoresistance in *SLFN11^−/−^* EWS. To assess the combined effect of both drugs, we employed BRAID modeling, which quantifies interaction strength through the kappa (κ) metric.^43,44^ Combination treatment with SN-38 and FSG67 significantly enhanced cell death in *SLFN11^−/−^* cells, demonstrating a modest synergistic effect (κ∼0.8). In contrast, no such synergy was observed in *SLFN11*-expressing WT cells.

In summary, these findings reveal a shift towards GPL biosynthesis associated with *SLFN11* knockout in EWS that contributes to therapeutic resistance and exposes metabolic vulnerability. The co-targeting DNA damage and lipid biosynthesis pathways elicits a synergistic anti-tumor effect and offers a promising strategy to overcome DDA resistance.

### Limitations of the study

While our study offers important insights into how *SLFN11* knockout drives metabolic adaptation and therapy resistance, several limitations should be acknowledged. *In vivo* validation was restricted to a single xenograft model (ES-8), which may not fully represent the metabolic heterogeneity across EWS tumors. Additionally, while transcriptomic and metabolomic data were integrated across multiple cell lines, the functional consequences of *GPD2* suppression and lipid remodeling were not evaluated using genetic rescue or knockdown models.

Moreover, isotope tracing was performed under nutrient-replete, *in vitro* conditions, which may not fully mimic the metabolic constraints present in the tumor microenvironment. While BRAID analysis revealed synergy between SN-38 and FSG67 in *SLFN11^−/−^* cells, the overall efficacy of this combination was modest, suggesting a need for further optimization or alternative targets. Finally, the regulatory mechanism linking *SLFN11* and *GPD2* remains correlative and requires deeper investigation to establish causality and therapeutic relevance.

## Resource Availability

### Lead contact

Further information and requests for resources and reagents should be directed to and will be fulfilled by the lead contact, Puneet Bagga.

### Materials availability

No unique reagents were generated in this study.

### Data and code availability

- All data reported in this study are available from the lead contact upon request.
- Any additional information required to reanalyze the data reported in this paper is available from the lead contact upon request.

## Acknowledgments

This study was supported by funding from the American Lebanese Syrian Associated Charities (ALSAC) and St. Jude Children’s Research Hospital (SJCRH). We thank the Hartwell Center and the Center for Applied Bioinformatics at SJCRH for generating and analyzing Transcriptomics data. Special thanks to Weixing Zhang of the Biomolecular Nuclear Magnetic Resonance Center. We also acknowledge the support of the Center for InVivo Imaging & Therapy and the Animal Resource Center at SJCRH. We would like to extend our sincere thanks to Cara Goodrum for tumor implantation, Ana Oliveira Souza, Kiel Neuman, and Spenser Simpson for their assistance, Laura Sanchez, and Sabah Nisar for their valuable help. Figures were created with BioRender.com under an academic license.

## Author Contributions

K.C. and P.B. conceived and designed the study. K.C., R.B., S.S., C.K., H.T., A.L., A.R., and W.Z. performed experiments. K.C., Z.F.Y., R.B., Y.W., X.W, Y.F., and A.S. analyzed the data. M.K., E.S. and M.A.D. provided the cell lines. K.C. wrote the manuscript with input from Z.F.Y., H.T., Y.W., Y.F., A.A.H., M.M., A.S., E.S., M.A.D. and P.B. P.B. supervised the study.

## Declaration of Interests

The authors declare no competing interests.

## Supplemental Information

Supplemental information can be found online at **“xxxx”**

## Methods

### Cell lines and culture

The cells used in this study were kindly provided by Dr. Elizabth Stewart of St. Jude Children’s Research Hospital. Both WT and *SLFN11*^−/−^ cell lines from a total of seven EWS cell lines—ES-8, EW-8, CADO ES-1, RD ES-1, and SK ES-1 were utilized in this study (Table S1). ES-8 and EW-8 cell lines were cultured in DMEM without glutamine with 10% fetal bovine serum (FBS) and 1 % penicillin-streptomycin-glutamine (PSG) and maintained at 37 °C in a humidified incubator with 5% CO₂. SK ES-1 cells were cultured in McCoy’s 5A medium supplemented with 15 % FBS and 1% Pen-strep, CADO ES-1 cells were cultured in RPMI 1640, supplemented with 20 % FBS and 1 % PSG and RD ES-1 cells were cultured in RPMI 1640, supplemented with 15 % FBS and 1 % PSG.

### Metabolite extraction

One million cells were seeded in 6-well plates and grown overnight. The cells were washed twice with ice cold 1x phosphate buffered saline (PBS), then 500 µL freezing 80% acetonitrile (LC-MS grade, Life Technologies Corp) was added to the cells. The cells were harvested into 1.5 mL tubes and lysed in the presence of glass beads by Bullet Blender (Next Advance) at 4 °C until the sample were homogenized. The lysate was centrifuged at 21,000 x g for 5 min and metabolite containing supernatant was split into two aliquots and dried by speedvac, respectively. Lipids were extracted in a similar way by freezing 100% isopropanol (LC-MS grade, Sigma-Aldrich) from 1 to 3 million cells and dried.

### Metabolome profiling by LC-MS/MS

One aliquot of metabolite sample was resuspended in 1% acetonitrile plus 0.1% trifluoroacetic acid (100 μL/million cells), 50 μL of the sample was desalted by Ultra-C18 Micro spin columns (Harvard apparatus) and eluted by 125 µL of 80% acetonitrile plus 0.1% trifluoroacetic acid followed by speedvac drying. The sample was then resuspended in 30 μL of 5% formic acid and 2 μL was analyzed by acidic pH reverse phase LC-MS/MS with a self-packed column (75 μm × 15 cm with 1.9 µm C18 resin from Dr. Maisch) coupled with a Q Exactive HF Orbitrap MS (Thermo Fisher) in positive ion mode. Metabolites were eluted within a 50 min gradient (mobile phase A: 0.2% formic acid in H_2_O; mobile phase B: 0.2% formic acid in acetonitrile; flow rate: 0.25 μL/min). Another aliquot of metabolite sample was resuspended in a solvent containing 45% isopropanol, 5% acetonitrile and 50% H_2_O (20 μL/million cells) and 3 µL was analyzed by a ZIC-HILIC column (150 × 2.1 mm, EMD Millipore) coupled with a Q Exactive HF Orbitrap MS (Thermo Fisher) in negative ion mode. Metabolites were eluted within a 45 min gradient (mobile phase A: 10 mM ammonium acetate in 90% acetonitrile, pH=8; mobile phase B: 10 mM ammonium acetate in 100% H2O, pH=8; flow rate: 0.1 mL/min). The mass spectrometry methods for both metabolomics analyses were set up with the following parameters: one MS1 scan (120,000 resolution, 100-1000 m/z, 3 x 10^6^ AGC and 50 ms maximal ion time) followed by 20 data-dependent MS2 scans (30,000 resolution, 2 x 10^5^ AGC, 45 ms maximal ion time, HCD, Stepped NCE (50, 100, 150), and 20 s dynamic exclusion).

Metabolomics mass spec data were converted into the mzXML format and processed using in-house JUMPm algorithm.^78^ Briefly, metabolite peak features were detected for each sample and aligned among all the compared samples. Metabolites were annotated by matching the retention time, accurate mass/charge ratio, and MS/MS fragmentation data to our in-house authentic compound library or matching to downloaded experimental MS/MS library (MoNA, https://mona.fiehnlab.ucdavis.edu/) by accurate mass/charge ratio and MS/MS spectrum. The dot product algorithm was employed to score the identifications. Peak intensities were used for metabolite quantification. The data was further normalized for batch effect removal (LIMMA R package,^79^ followed by quantile normalization.^80^ Quantification and statistical analysis were done by calculating fold changes and P values between different groups using the LIMMA R package. MetaboAnalyst6.0 (www.metaboanalyst.ca)^38,39,81^ was used to generate metabolite set enrichment analysis (MSEA) plots for pathway analysis.

### *In vitro* stable isotope tracing analysis by LC-MS/MS

One million cells were seeded and allowed to adhere overnight for each experiment. For the isotope tracing experiment, cells were washed with PBS and subsequently incubated with a medium containing ^13^C isotope-labeled tracer [either of U-^13^C glucose (CLM-1396; 99%) or U-^13^C acetate (CLM-440; 99%)] of interest supplemented with 5% dialyzed FBS (Gibco:A33820-01). The medium was adjusted to include glucose at a concentration of 10 mM,^59^ glutamine at 4 mM,^59^ acetate at 0.4 mM.^42^ Cells were incubated under these conditions for 24 h. After incubation, cells were washed twice with ice-cold 1x PBS to remove residual medium. Metabolite extraction and LC-MS/MS methods were the same as untargeted metabolome profiling analyses except positive ion mode analysis for metabolomics was not pursued based on the metabolite targets of interested.

MS raw data was converted into the mzXML format and processed using in-house JUMPm algorithm. Briefly, the feature peaks of the isotopologues of all target metabolites were extracted and aligned among all the compared samples followed by natural isotope abundance correction and tracer impurity correction. The identified metabolites were validated by comparing with authentic standard compounds at the following parameters: mass/charge ratio, LC retention time and MS/MS spectra. The peak intensities were used for quantification and calculating the labeling percentage of the isotopologues for each target metabolite.

### Xenograft studies in mice

Athymic nude immunodeficient mice were obtained from Charles River Laboratories. All animal experiments were conducted in accordance with protocols approved by the Institutional Animal Care and Use Committee (IACUC), and efforts were made to minimize animal suffering. Mice were housed under a 12-h light/dark cycle and provided food and water ad libitum. To generate EWS orthotopic xenografts, ES-8 WT and ES-8 *SLFN11*^−/−^ cells were suspended in Matrigel (Corning, Catalogue no: 356234) at a concentration of 20,000 cells/μL. The cell suspensions were injected into the bone marrow as previously described in Stewart et al.^3^, using a Hamilton syringe fitted with a 25-gauge needle. Once tumors became palpable and reached a diameter of approximately 20 mm, mice were euthanized and the tumors grown surrounding the femur were harvested and used for downstream analyses.

### Preparation for tumor extracts

Metabolite extraction from tumor extracts was done following the method described in Patel *et al.*^82^ Briefly, frozen tissue (150–200 mg) from tumor grown surrounding the femur was ground with 0.1 M HCl/methanol (2:1 vol wt) at 40°C followed by extraction with ice-cold ethanol. The supernatant was clarified by centrifugation, lyophilized, and resuspended in 500 µL of phosphate-buffered (25 mM, pH-7) D_2_O solution containing 3-(trimethylsilyl) propionic-2,2,3,3-d_4_ acid (0.05 wt%).

### NMR spectroscopy of tumor extracts for ^1^H NMR

¹H NMR spectra of tumor extracts were acquired at 298 K on a 600 MHz Bruker Avance NEO NMR spectrometer (Bruker BioSpin, Billerica, MA) equipped with a 5 mm TCI cryoprobe. Acquisition parameters included a spectral width of 13.0 ppm, 16k data points, a relaxation delay of 2 s, an acquisition time of 1.1 s, and 64 scans. The residual H₂O signal was suppressed using the excitation sculpting technique.^83^ Free induction decays (FIDs) were Fourier-transformed and analyzed using Bruker Topspin 4.3.0 software. The peak intensity of different metabolites was measured, and choline-containing compounds were identified based on their characteristic chemical shifts: phosphocholine (PCh, 3.226 ppm), glycerophosphocholine (GPC, 3.235 ppm), and free choline (Cho, 3.208 ppm).^49^

### RNA extraction, sequencing, and data analysis

RNA was extracted from *in vitro* samples using the RNeasy Mini Kit (QIAGEN, Cat. No. 74104). Expression profiles were generated from biological triplicates, as well as biological duplicates of all cell lines. RNA sample quality was assessed using the Agilent 2200 TapeStation System, and data analysis was performed with TapeStation Analysis Software 5.1. RNA-Seq libraries were prepared using the TruSeq Total RNA protocol and sequenced in pools of 5 to 7 samples per lane on a V3 flow cell (HiSeq 2000/2500). Where necessary, additional sequencing (“top-off”) was performed in rapid mode on the HiSeq 2500 to ensure data analysis was completed within a clinically appropriate time frame.^84^ The RNA seq reads were mapped to mouse genome (gencode M22) using STAR2.7. After mapping, the gene count matrix for each sample was generated by RSEM. The differentially expressed genes were identified using limma package with count matrix as input. The cutoff for differential gene is log2fold change > 0.5 and an FDR-adjusted p ≤ 0.05. Boxplots and volcano plots were generated using in-house scripts.

### Immunoblotting

The following antibodies were used: *SLFN11* (Sigma: HPA023030), *GPD2* (Proteintech: 17219-1-AP), β-actin (Cell Signaling Technology: 4970), and secondary antibody (Cell Signaling Technology: 7074P2). Cells were lysed using 1x-lysis buffer (Cell Signaling Technology: 9803), vortexed, and centrifuged at 4 °C for 15 mins. The supernatant was collected, and protein concentration was measured using the BCA Protein Quantification Assay (Thermo: 23228). Proteins were separated by SDS-PAGE and transferred to PVDF membranes (Merck: ISEQ00005). Membranes were blocked with 5% skimmed milk or BSA for 1 h at room temperature. After blocking, the PVDF membrane was incubated with primary antibodies at 4 °C overnight on a rocker. Secondary antibodies were diluted in TBST and added to the membranes, followed by incubation for 2 h at room temperature. Protein bands were visualized using the ChemiDocTM Touch Imaging System.

### High-throughput drug combination screening and BRAID analysis

ES-8 WT and ES-8 *SLFN11*^−/−^ cells were seeded in 384-well plates with an automatic dispenser (WellMate, Thermo Scientific) at a density of 3000 cells/well. Following 24 h of incubation, concentrated compounds were pin transferred in the 30uL cell suspension using the Biomek FX Laboratory Automation Workstation (Beckman Coulter, Inc) resulting in a 1:1000 dilution. DMSO was used as negative control. A BRAID-format drug combination screen was performed using SN-38 as the anchor drug. SN-38 was assessed across seven concentrations in combination with ten concentrations of each of three partner compounds: FSG67, TH9619, CB-839, and SN-38 (self-control). Source plate stock concentrations were prepared at 1,000× for a final assay dilution of: SN-38 (1 μM-0.05 nM) and FSG67 (200 μM-10 nM). Cells were treated in triplicate technical replicates across two biological replicates. After 72 h, cell viability was measured using CellTiter-Glo (Promega: G7573), and luminescence was recorded using an EnVision plate reader (PerkinElmer). Dose-response surfaces were modeled using the BRAID (Bivariate Response to Additive Interacting Doses) approach to quantify drug interactions.^43,44^ The κ (kappa) parameter indicates interaction type, where κ > 0 denotes synergy, κ = 0 indicates additivity, and κ < 0 indicates antagonism. IAE50 and IAE90 were calculated to assess the overall efficacy of each combination at 50% and 90% inhibition thresholds, respectively. IDMA and IDMB correspond to the EC50 values of the partner drug and the anchor drug, respectively. Statistical significance of interaction terms was evaluated based on 95% confidence intervals for κ.

### Kaplan-Meier survival analysis protocol for gene expression data

Kaplan Meier survival analysis was performed by categorizing patients into high- and low-expression groups using the “minimum p-value approach” as the cutoff for each gene. This method iteratively applies ascending gene expression values as thresholds to divide the cohort into two groups and evaluates the p-value at each step using a log-rank test. The optimal cutoff value was determined and used to generate the corresponding Kaplan-Meier survival curves. The analysis was conducted on the R2: Genomics Analysis and Visualization Platform (http://r2.amc.nl)^30^ using survival and gene expression data from the GSE17679 dataset.^37^

### mRNA expression profiling

mRNA expression profile for *SLFN11* was analyzed across multiple cancer types using the Cancer Cell Line Encyclopedia (CCLE) database.^28^ The analysis included cell lines representing EWS, small cell lung cancer, ovarian cancer, prostate cancer, gastric cancer, breast cancer, colorectal cancer, acute myeloid leukemia, neuroblastoma, renal cell carcinoma, glioblastoma, and hepatocellular carcinoma.

### DepMap-gene dependency analysis

CRISPR-based gene dependency data were obtained from the DepMap database (https://depmap.org/portal) (<u>DepMap version:</u> <u>24Q4</u>), which was normalized using the Chronos algorithm. Dependency scores represent the likelihood of a gene being essential for cell survival, with lower scores indicating higher dependency. Genes with dependency scores below [threshold, e.g., −0.5] were considered essential. A *t*-test was used to compare the dependency score of *SLFN11* across each cancer lineage or primary disease type against all the cell lines. P-values were adjusted using the Benjamini-Hochberg method for false discovery rate (FDR) correction.

### Correlation analysis of gene expression

Transcriptomic data were obtained from the Ewing Sarcoma Cell Line Atlas (ESCLA)^30^ and the inflammatory gene profiling dataset of the Ewing sarcoma family of tumors.^37^ Batch effects due to treatment conditions, cell lines, or diagnostic subtypes were removed using the limma package (version 3.60.4).^79^ Pearson correlation coefficients were calculated on log2-normalized expression values, and the significance of correlations was tested using the cor.test function in R.

### Statistical analysis

MetaboAnalyst 6.0, R software and MATLAB (R2023b) were used to conduct statistical analysis of metabolomics data. For additional data analysis GraphPad Prism V10.4.0 was used to conduct Student’s t test. Data visualization used GraphPad Prism, MATLAB, and R software. Descriptions of individual statistical analyses can be found in the figure legends.

## Abbreviations

EWS: ewing sarcoma
*SLFN11*: schlafen family member 11
DDA: DNA-damaging agent
GPL: glycerophospholipid
*GPAT1*: glycerol-3-phosphate acyltransferase 1
*GPD2*: glycerol-3-phosphate dehydrogenase 2
*EWSR1*: ewing sarcoma breakpoint region 1
ETS: erythroblast transformation specific
*FLI1*: Friend Leukemia Integration 1
PARP: poly (ADP-ribose) polymerase
G3P: glycerol-3-phosphate
CCLE: Cancer Cell Line Encyclopedia
DepMap: cancer dependency map
CRISPR: clustered regularly interspaced short palindromic repeats
RNAi: RNA interference
FDR: false discovery rate
TCGA: The Cancer Genome Atlas Program
ESCLA: Ewing Sarcoma Cell Line Atlas
RNA-seq: RNA sequencing
PCA: principal component analysis
PC1: first principal component
Log_2_FC: Log_2_ fold change
WT: wild type
*SLFN11^−/−^*: *SLFN11* knock out
Pearson R: Pearson correlation coefficient
G3PS: glycerol-3-phosphate shuttle
DHAP: dihydroxyacetone phosphate
ETC: electron transport chain
LC/MS: Liquid chromatography–mass spectrometry
PE: phosphatidylethanolamine
PC: phosphatidylcholine
PI: phosphatidylinositol
PG: phosphatidylglycerol
PA: phosphatidic acid
DAG: diacylglycerol
*AGPAT4*: 1-Acylglycerol-3-Phosphate O-Acyltransferase 4
MUFA: monounsaturated fatty acid
^1^H NMR: proton nuclear magnetic resonance
PCh: phosphocholine
GPC: glycerophosphocholine
BRAID: Bivariate Response to Additive Interacting Doses
*GPAT1*: glycerol-3-phosphate acyltransferase 1
IAE: Index of Achievable Efficacy
IDMA: inhibitory dose for the anchor drug
IDMB: inhibitory dose for the partner drug
EC50: half maximal effective concentration
RPA1: replication protein A
MCM3: minichromosome maintenance complex component 3
CDC45: cell division cycle 45
PCNA: proliferating cell nuclear antigen
ATR: Ataxia Telangiectasia and Rad3-related
CDT: chromatin licensing and DNA replication factor 1
DDB1: damage-binding protein 1
CUL4: cullin 4
NSCLC: non-small cell lung cancer
ccRCC: clear cell renal cell carcinoma
mTOR: mammalian target of rapamycin
FASN: fatty acid synthase
ACC: acetyl-CoA carboxylase
SCD1: stearoyl-CoA desaturase 1.

## Notes

### Competing Interest Statement

The authors have declared no competing interest.

