## Supplemental Figures and Tables for "*SLFN11* Loss-Induced Chemoresistance is Associated with Overexpression of Glycerophospholipid Biosynthesis in Ewing Sarcoma"

**Table S1.** Summary of EWS cell lines used in this study.

| Cell Line | Source | Age (years) | Sex | Primary Site | Metastatic Site | Translocation Type |
|---|---|---|---|---|---|---|
| ES-8 | SJCRH | 10 | M | Left proximal humerus | Femur | EWSR1 Exon 7 to FLI1 Exon 5 (Type II) |
| SK ES-1 | ATCC | 18 | M | Bone | - | EWSR1 Exon 7 to FLI1 Exon 5 (Type II) |
| EW-8 | SJCRH | 17 | M | Abdominal mass | - | EWSR1 Exon 7 to FLI1 Exon 6 (Type I) |
| RD ES-1 | ATCC | 19 | M | Bone | - | EWSR1 Exon 7 to FLI1 Exon 5 (Type II) |
| CADO ES-1 | Center for Adult Diseases Osaka | 19 | F | | Lung | EWSR1-ERG |

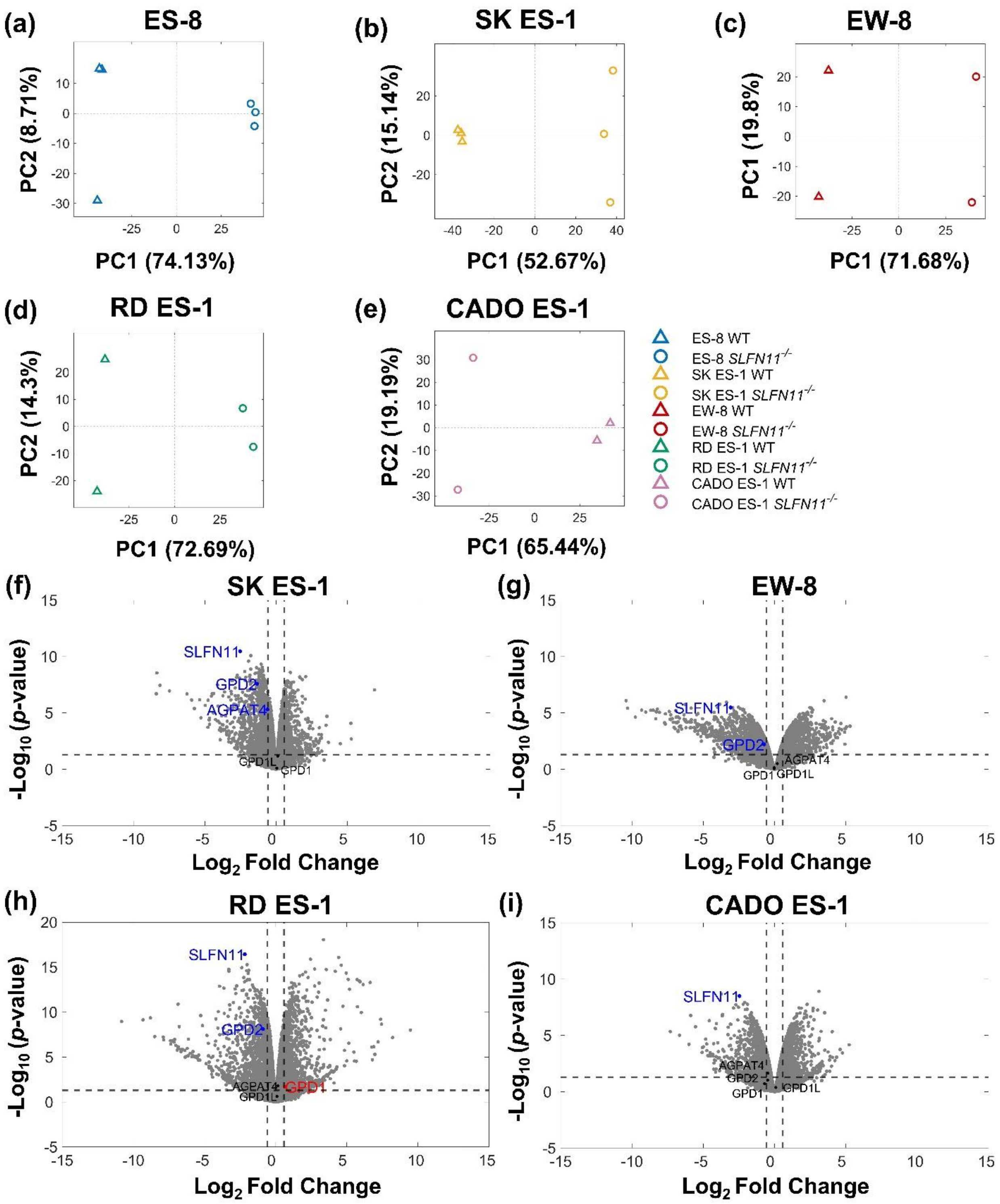

(a)
ES-8
PC2 (8.71%)
PC1 (74.13%)
(b)
SK ES-1
PC2 (15.14%)
PC1 (52.67%)
(c)
EW-8
PC1 (19.8%)
PC1 (71.68%)
(d)
RD ES-1
PC2 (14.3%)
PC1 (72.69%)
(e)
CADO ES-1
PC2 (19.19%)
PC1 (65.44%)
ES-8 WT
ES-8 SLFN11-/-
SK ES-1 WT
SK ES-1 SLFN11-/-
EW-8 WT
EW-8 SLFN11-/-
RD ES-1 WT
RD ES-1 SLFN11-/-
CADO ES-1 WT
CADO ES-1 SLFN11-/-
(f)
SK ES-1
SLFN11
GPD2
AGPAT4
GPD1L
GPD1
(g)
EW-8
SLFN11
GPD2
AGPAT4
GPD1
GPD1L
(h)
RD ES-1
SLFN11
GPD2
AGPAT4
GPD1
GPD1L
(i)
CADO ES-1
SLFN11
AGPAT4
GPD2
GPD1
GPD1L
-Log10 (p-value)
Log2 Fold Change

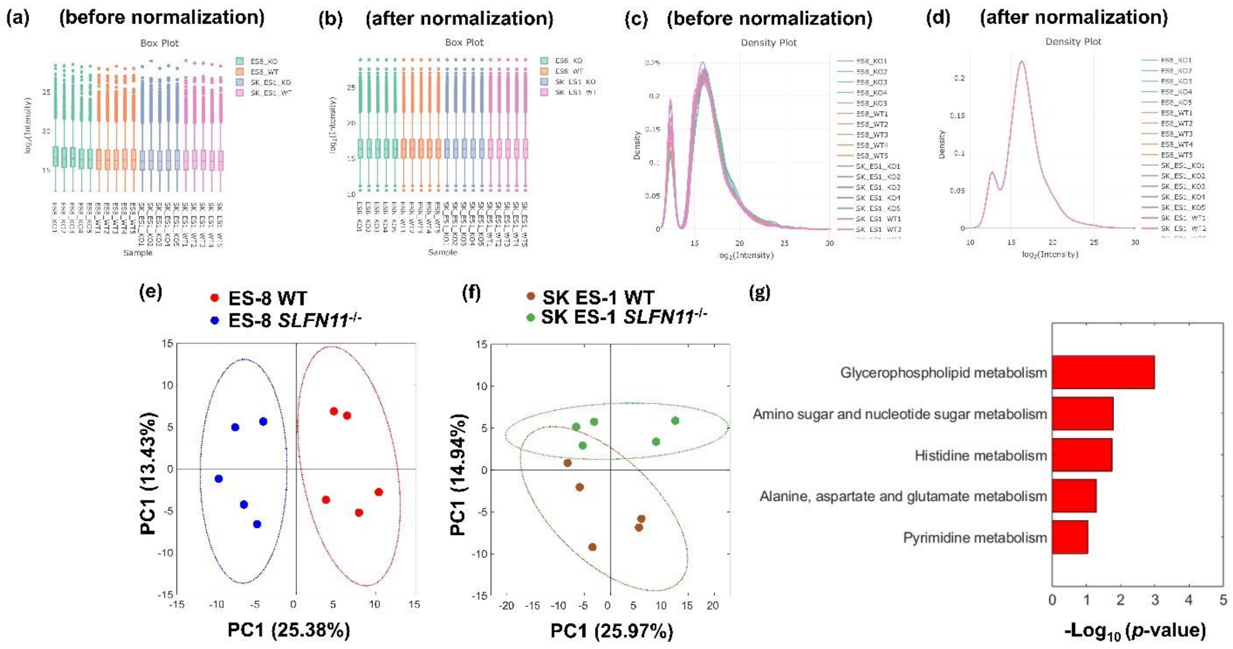

(a) (before normalization)
Box Plot
(b) (after normalization)
Box Plot
(c) (before normalization)
Density Plot
(d) (after normalization)
Density Plot
(e)
ES-8 WT
ES-8 SLFN11-/-
PC1 (13.43%)
PC1 (25.38%)
(f)
SK ES-1 WT
SK ES-1 SLFN11-/-
PC1 (14.94%)
PC1 (25.97%)
(g)
Glycerophospholipid metabolism
Amino sugar and nucleotide sugar metabolism
Histidine metabolism
Alanine, aspartate and glutamate metabolism
Pyrimidine metabolism
-Log10 (p-value)

**Table S2.** Upregulated metabolites in ES-8 *SLFN11*$^{-/-}$ cell line compared to ES-8 WT.

| **Metabolite** | **HMDB ID** | **FDR** | **Log2Fold (ES-8 SLFN11$^{-/-}$vs. ES-1 WT)** |
|---|---|---|---|
| Dihydroorotate | HMDB0003349 | 2.43E-03 | 0.51 |
| Uridine 5'-Diphosphate | HMDB0000295 | 1.52E-01 | 0.54 |
| Cytidine Monophosphate | HMDB0000095 | 7.33E-03 | 0.7 |
| Adenosine 5'-Triphosphate | HMDB0000538 | 1.69E-01 | 0.77 |
| D-Ribose 5-Phosphate | HMDB0001548 | 2.29E-03 | 0.91 |
| Heptadecanoic acid | HMDB0002259 | 1.18E-02 | 1.05 |
| Xanthine | HMDB0000292 | 9.61E-09 | 1.07 |
| Folate | HMDB0000121 | 9.98E-05 | 1.12 |
| Vaccenic Acid | HMDB0240219 | 1.25E-04 | 1.2 |
| Uridine 5'-Triphosphate | HMDB0000285 | 4.78E-02 | 1.29 |
| Serine | HMDB0000187 | 1.61E-01 | 1.31 |
| Nervonic Acid | HMDB0002368 | 1.31E-04 | 1.4 |
| Phosphoenolpyruvate | HMDB0000263 | 9.31E-02 | 1.61 |
| Uridine Monophosphate | HMDB0000288 | 6.61E-09 | 2.22 |
| **Glycerol 3-Phosphate** | **HMDB0000126** | **1.43E-06** | **2.28** |
| Inosine | HMDB0000195 | 4.32E-07 | 3.37 |
| Citrate | HMDB0000094 | 6.80E-02 | 3.65 |
| **O-Phosphoethanolamine** | **HMDB0000224** | **9.07E-10** | **3.9** |
| Inosine 5'-Monophosphate | HMDB0000175 | 2.50E-04 | 4.01 |

**Table S3.** Upregulated metabolites in SK ES-1 *SLFN11*$^{-/-}$ cell line compared to SK ES-1 WT.

| Metabolite | HMDB ID | FDR | Log2Fold (SK ES-1 SLFN11$^{-/-}$ vs.SK ES-1 WT) |
|---|---|---|---|
| Cytidine Monophosphate | HMDB0000095 | 4.13E-03 | 0.8 |
| Uridine Monophosphate | HMDB0000288 | 1.29E-04 | 1.24 |
| **Glycerol 3-Phosphate** | **HMDB0000126** | **7.33E-04** | **1.5** |
| L-Glutamic Acid | HMDB0000148 | 3.07E-01 | 1.93 |
| **O-Phosphoethanolamine** | **HMDB0000224** | **1.15E-08** | **3.63** |

**Table S4.** Isotopologue distribution pattern of metabolites labeled with U-$^{13}$C glucose in ES-8 cell line.

| Labeling % | Isotopologue | ES-8 WT | ES-8 WT | ES-8 WT | ES-8 *SLFN11$^{-/-}$* | ES-8 *SLFN11$^{-/-}$* | ES-8 *SLFN11$^{-/-}$* |
|---|---|---|---|---|---|---|---|
| Pyruvate | M+3 | 99.86 | 99.94 | 100.19 | 99.09 | 89.72 | 98.16 |
| Lactate | M+3 | 87.68 | 90.69 | 90.48 | 91.14 | 91.18 | 91.27 |
| Citrate | M+2 | 32.57 | 30.29 | 42.51 | 32.96 | 34.85 | 34.44 |
| Glutamate | M+2 | 23.89 | 22.43 | 23.20 | 22.06 | 21.18 | 21.80 |
| Fumarate | M+2 | 23.15 | 21.71 | 19.81 | 25.12 | 25.92 | 24.39 |
| Malate | M+2 | 25.24 | 24.47 | 28.99 | 25.13 | 26.96 | 25.65 |
| Aspartate | M+2 | 20.41 | 22.32 | 19.76 | 16.11 | 15.10 | 15.65 |
| Aspartate | M+3 | 17.25 | 20.49 | 15.14 | 12.23 | 12.91 | 12.30 |
| Malate | M+3 | 21.09 | 23.41 | 22.95 | 22.36 | 25.42 | 23.10 |
| Fumarate | M+3 | 16.64 | 23.03 | 14.62 | 20.67 | 21.14 | 21.71 |
| G3P | M+3 | 35.34 | 51.64 | 39.74 | 89.67 | 92.66 | 92.59 |
| DHAP | M+3 | 90.29 | 92.20 | 89.54 | 92.64 | 93.98 | 95.30 |
| F6P | M+3 | 0.09 | 0.07 | 0.11 | 0.12 | 0.09 | 0.06 |
| F6P | M+6 | 48.64 | 59.30 | 37.00 | 35.63 | 27.38 | 31.30 |

**Table S5.** Isotopologue distribution pattern of metabolites labeled with U-$^{13}$C glucose in SK ES-1 cell line.

| Labeling % | Isotopologue | SK ES-1 WT | SK ES-1 WT | SK ES-1 WT | SK ES-1 SLFN11$^{-/-}$ | SK ES-1 SLFN11$^{-/-}$ | SK ES-1 SLFN11$^{-/-}$ |
|---|---|---|---|---|---|---|---|
| Lactate | M+3 | 78.39 | 76.30 | 83.01 | 73.24 | 89.68 | 89.17 |
| Citrate | M+2 | 22.89 | 36.26 | 39.81 | 30.14 | 31.77 | 34.87 |
| Glutamate | M+2 | 23.73 | 21.50 | 23.16 | 21.14 | 21.83 | 22.17 |
| Fumarate | M+2 | 17.91 | 16.01 | 19.15 | 15.86 | 15.46 | 18.73 |
| Malate | M+2 | 20.37 | 17.93 | 18.94 | 16.61 | 17.78 | 18.83 |
| Aspartate | M+2 | 18.82 | 6.94 | 15.27 | 9.96 | 15.71 | 15.76 |
| Aspartate | M+3 | 6.58 | 3.17 | 4.75 | 3.76 | 4.89 | 5.94 |
| Malate | M+3 | 10.46 | 9.29 | 9.71 | 7.53 | 8.37 | 8.82 |
| Fumarate | M+3 | 4.57 | 3.99 | 4.68 | 3.26 | 3.23 | 5.21 |
| G3P | M+3 | 89.06 | 89.74 | 89.49 | 94.42 | 95.13 | 93.40 |
| DHAP | M+3 | 93.41 | 89.88 | 89.01 | 93.22 | 94.81 | 93.61 |
| F6P | M+3 | 5.58 | 3.91 | 4.11 | 3.76 | 2.98 | 3.23 |
| F6P | M+6 | 76.54 | 83.01 | 85.29 | 85.11 | 87.78 | 91.21 |

**Table S6.** Up and downregulated pathways in ES-8 *SLFN11*$^{-/-}$ cell line compared to ES-8 WT.

| **Upregulated pathway** | **Overlapped HMDB IDs** | **Overlapped metabolites** | **# metabolites overlapped** | **# metabolites in the pathway** | **p-value** |
|---|---|---|---|---|---|
| Pyrimidine metabolism | HMDB0000295 | Uridine 5'-diphosphate | 6 | 39 | 8.26E-05 |
| | HMDB0000288 | Uridine 5'-monophosphate | | | |
| | HMDB0000095 | Cytidine monophosphate | | | |
| | HMDB0001202 | dCMP | | | |
| | HMDB0000528 | 4,5-Dihydroorotic acid | | | |
| | HMDB0000828 | Ureidosuccinic acid | | | |
| Purine metabolism | HMDB0001548 | D-Ribose 5-phosphate | 7 | 70 | 3.28E-04 |
| | HMDB0000462 | Allantoin | | | |
| | HMDB0000538 | Adenosine triphosphate | | | |
| | HMDB0000175 | Inosinic acid | | | |
| | HMDB0000195 | Inosine | | | |
| | HMDB0000292 | Xanthine | | | |
| | HMDB0000299 | Xanthosine | | | |
| One carbon pool by folate | HMDB0000121 | Folic acid | 4 | 26 | 1.47E-03 |
| | HMDB0000187 | Serine | | | |
| | HMDB0000538 | Adenosine triphosphate | | | |
| | HMDB0000939 | S-Adenosylhomocysteine | | | |
| Alanine, aspartate, and glutamate metabolism | HMDB0000094 | Citric acid | 3 | 28 | 1.72E-02 |
| | HMDB0000828 | Ureidosuccinic acid | | | |
| | HMDB0001254 | Glucosamine 6-phosphate | | | |
| Amino sugar and nucleotide sugar metabolism | HMDB0001254 | Glucosamine 6-phosphate | 3 | 42 | 4.98E-02 |
| | HMDB0000290 | Uridine diphosphate-N-acetylglucosamine | | | |
| | HMDB0001163 | Guanosine diphosphate mannose | | | |
| Glycerophospholipid metabolism | HMDB0000126 | Glycerol 3-phosphate | 2 | 36 | 1.62E-01 |
| | HMDB0000224 | O-Phosphoethanolamine | | | |
| **Downregulated pathway** | **Overlapped HMDB IDs** | **Overlapped metabolites** | **# metabolites overlapped** | **# metabolites in the pathway** | **p-value** |
| Alanine, aspartate, and glutamate metabolism | HMDB0000812 | N-Acetyl-L-aspartic acid | 5 | 28 | 1.08E-04 |
| | HMDB0006483 | D-Aspartic acid | | | |
| | HMDB0000168 | L-Asparagine | | | |

| | | | | | |
|---|---|---|---|---|---|
| | HMDB0000208 | Oxoglutaric acid | | | |
| | HMDB0000134 | Fumaric acid | | | |
| Citrate cycle (TCA cycle) | HMDB0000156 | Malic acid | 4 | 20 | 3.67E-04 |
| | HMDB0000134 | Fumaric acid | | | |
| | HMDB0000072 | cis-Aconitic acid | | | |
| | HMDB0000208 | Oxoglutaric acid | | | |
| Pyruvate metabolism | HMDB0001066 | S-Lactoylglutathione | 3 | 23 | 7.76E-03 |
| | HMDB0000156 | Malic acid | | | |
| | HMDB0000134 | Fumaric acid | | | |
| Arginine biosynthesis | HMDB0000208 | Oxoglutaric acid | 2 | 14 | 2.60E-02 |
| | HMDB0000134 | Fumaric acid | | | |
| Histidine metabolism | HMDB0000033 | Carnosine | 2 | 16 | 3.35E-02 |
| | HMDB0000001 | 1-Methylhistidine | | | |

**Table S7.** Up and downregulated pathways in SK ES-1 *SLFN11*$^{-/-}$ cell line compared to SK ES-1 WT.

| **Upregulated pathway** | **Overlapped HMDB IDs** | **Overlapped metabolites** | **# metabolites overlapped** | **# metabolites in the pathway** | **p-value** |
|---|---|---|---|---|---|
| Glycerophospholipid metabolism | HMDB0000126 | Glycerol 3-phosphate | 3 | 36 | 9.30E-03 |
| | HMDB0000224 | O-Phosphoethanolamine | | | |
| | HMDB0008834 | PE(14:0/20:1(11Z)) | | | |
| Amino sugar and nucleotide sugar metabolism | HMDB0001254 | Glucosamine 6-phosphate | 3 | 42 | 1.42E-02 |
| | HMDB0000290 | Uridine diphosphate-N-acetylglucosamine | | | |
| | HMDB0001163 | Guanosine diphosphate mannose | | | |
| Histidine metabolism | HMDB0000033 | Carnosine | 2 | 16 | 1.62E-02 |
| | HMDB0000148 | Glutamic acid | | | |
| Glutathione metabolism | HMDB0000148 | Glutamic acid | 2 | 28 | 4.66E-02 |
| | HMDB0001049 | gamma-Glutamylcysteine | | | |
| Alanine, aspartate and glutamate metabolism | HMDB0000148 | Glutamic acid | 2 | 28 | 4.66E-02 |
| | HMDB0001254 | Glucosamine 6-phosphate | | | |
| Pyrimidine metabolism | HMDB0000288 | Uridine 5'-monophosphate | 2 | 39 | 8.42E-02 |
| | HMDB0000095 | Cytidine monophosphate | | | |
| **Downregulated pathway** | **Overlapped HMDB IDs** | **Overlapped metabolites** | **# metabolites overlapped** | **# metabolites in the pathway** | **p-value** |
| Alanine, aspartate, and glutamate metabolism | HMDB0006483 | D-Aspartic acid | 5 | 28 | 3.91E-05 |
| | HMDB0000828 | Ureidosuccinic acid | | | |
| | HMDB0000208 | Oxoglutaric acid | | | |
| | HMDB0000134 | Fumaric acid | | | |
| | HMDB0000254 | Succinic acid | | | |
| Citrate cycle (TCA cycle) | HMDB0000156 | Malic acid | 4 | 20 | 1.64E-04 |
| | HMDB0000254 | Succinic acid | | | |
| | HMDB0000134 | Fumaric acid | | | |
| | HMDB0000208 | Oxoglutaric acid | | | |

| | | | | | |
|---|---|---|---|---|---|
| Purine metabolism | HMDB0001517 | AICAR | 6 | 70 | 4.19E-04 |
| | HMDB0000133 | Guanosine | | | |
| | HMDB0000045 | Adenosine monophosphate | | | |
| | HMDB0000034 | Adenine | | | |
| | HMDB0000195 | Inosine | | | |
| | HMDB0000292 | Xanthine | | | |
| One carbon pool by folate | HMDB0000121 | Folic acid | 4 | 26 | 4.77E-04 |
| | HMDB0000187 | Serine | | | |
| | HMDB0000092 | Dimethylglycine | | | |
| | HMDB0000099 | L-Cystathionine | | | |
| Arginine biosynthesis | HMDB0000208 | Oxoglutaric acid | 3 | 14 | 9.84E-04 |
| | HMDB0000904 | Citrulline | | | |
| | HMDB0000134 | Fumaric acid | | | |
| Glycine, serine, and threonine metabolism | HMDB0000092 | Dimethylglycine | 4 | 33 | 1.21E-03 |
| | HMDB0001149 | 5-Aminolevulinic acid | | | |
| | HMDB0000099 | L-Cystathionine | | | |
| | HMDB0000187 | Serine | | | |
| Pyruvate metabolism | HMDB0001066 | S-Lactoylglutathione | 3 | 23 | 4.37E-03 |
| | HMDB0000156 | Malic acid | | | |
| | HMDB0000134 | Fumaric acid | | | |
| D-Amino acid metabolism | HMDB0006483 | D-Aspartic acid | 2 | 15 | 2.02E-02 |
| | HMDB0000187 | Serine | | | |
| Butanoate metabolism | HMDB0000208 | Oxoglutaric acid | 2 | 15 | 2.02E-02 |
| | HMDB0000254 | Succinic acid | | | |

**Table S8.** Isotopologue distribution pattern of metabolites labeled with U$^{13}$C acetate in ES-8 cell line.

| Phospholipid species | Isotopologues | % Enrichment | | | |
|---|---|---|---|---|---|
| | | **ES-8 WT** | | **ES-8 *SLFN11*$^{-/-}$** | |
| | | **Avg** | **SD** | **Avg** | **SD** |
| PE (34:1) | M+0 | 24.6 | 13.7 | 18.0 | 12.0 |
| | M+1 | 0.3 | 0.3 | 0.4 | 0.2 |
| | M+2 | 21.2 | 11.9 | 15.3 | 10.1 |
| | M+3 | 0.6 | 0.5 | 0.6 | 0.1 |
| | M+4 | 15.2 | 8.5 | 11.1 | 7.4 |
| | M+5 | 0.5 | 0.5 | 0.5 | 0.1 |
| | M+6 | 7.9 | 4.3 | 5.7 | 3.6 |
| | M+7 | 0.1 | 0.1 | 0.2 | 0.1 |
| | M+8 | 3.7 | 2.0 | 2.7 | 1.7 |
| | M+9 | 0.0 | 0.0 | 0.0 | 0.0 |
| | M+10 | 1.4 | 0.7 | 1.1 | 0.6 |
| | M+11 | 0.0 | 0.0 | 0.1 | 0.0 |
| | M+12 | 0.6 | 0.2 | 0.4 | 0.2 |
| PC (34:1) | M+0 | 16.8 | 11.4 | 14.1 | 2.7 |
| | M+1 | 0.3 | 0.1 | 0.2 | 0.1 |
| | M+2 | 19.1 | 13.2 | 16.2 | 2.9 |
| | M+3 | 1.0 | 0.6 | 0.8 | 0.2 |
| | M+4 | 15.0 | 10.4 | 12.7 | 2.3 |
| | M+5 | 0.5 | 0.2 | 0.4 | 0.2 |
| | M+6 | 8.4 | 5.6 | 7.0 | 1.4 |
| | M+7 | 0.5 | 0.3 | 0.4 | 0.1 |
| | M+8 | 3.7 | 2.4 | 3.0 | 0.6 |
| | M+9 | 0.2 | 0.1 | 0.1 | 0.0 |
| | M+10 | 1.4 | 0.9 | 1.2 | 0.2 |
| | M+11 | 0.1 | 0.0 | 0.1 | 0.0 |
| | M+12 | 0.5 | 0.3 | 0.4 | 0.1 |
| PG (34:1) | M+0 | 17.9 | 10.1 | 13.0 | 8.7 |
| | M+1 | 0.5 | 0.3 | 0.4 | 0.1 |
| | M+2 | 20.5 | 11.2 | 15.0 | 9.8 |
| | M+3 | 0.4 | 0.2 | 0.4 | 0.1 |
| | M+4 | 15.8 | 8.8 | 11.6 | 7.7 |
| | M+5 | 0.6 | 0.3 | 0.5 | 0.1 |
| | M+6 | 9.6 | 5.1 | 6.8 | 4.3 |
| | M+7 | 0.3 | 0.3 | 0.4 | 0.1 |
| | M+8 | 4.9 | 2.5 | 3.5 | 2.1 |
| | M+9 | 0.2 | 0.1 | 0.2 | 0.1 |

| | | | | | |
|---|---|---|---|---|---|
| | M+10 | 1.8 | 0.9 | 1.3 | 0.7 |
| | M+11 | 0.0 | 0.0 | 0.1 | 0.0 |
| | M+12 | 2.1 | 1.1 | 1.6 | 0.8 |
| PI (34:1) | M+0 | 19.3 | 10.6 | 14.2 | 9.4 |
| | M+1 | 0.6 | 0.4 | 0.5 | 0.1 |
| | M+2 | 21.5 | 11.9 | 15.8 | 10.5 |
| | M+3 | 0.3 | 0.3 | 0.3 | 0.0 |
| | M+4 | 17.3 | 9.8 | 12.6 | 8.5 |
| | M+5 | 0.3 | 0.1 | 0.2 | 0.1 |
| | M+6 | 9.5 | 5.2 | 6.8 | 4.5 |
| | M+7 | 0.6 | 0.5 | 0.5 | 0.1 |
| | M+8 | 4.5 | 2.2 | 3.2 | 1.8 |
| | M+9 | 0.2 | 0.1 | 0.2 | 0.1 |
| | M+10 | 1.6 | 0.8 | 1.1 | 0.7 |
| | M+11 | 0.0 | 0.0 | 0.0 | 0.0 |
| | M+12 | 0.7 | 0.3 | 0.5 | 0.2 |

**Table S9.** BRAID modeling results for SN-38-based drug combinations in ES-8 WT and *SLFN11*$^{-/-}$ cell lines. IDMA and IDMB represent the EC50 values of the partner drug (FSG67) and the anchor drug (SN-38), respectively. The synergy coefficient ($\kappa$) quantifies drug interaction, with positive values indicating synergy and negative values indicating antagonism. Near zero values indicate additive effect. $Kappa_{lo}$ and $Kappa_{hi}$ indicate the lower and upper bounds of the 95% confidence interval for the kappa ($\kappa$) interaction coefficient. IAE50 and IAE90 measure the combinatorial efficacy required to achieve 50% and 90% cell death, respectively. Cell line data are color-coded by metric type: pink indicates antagonistic $\kappa$ values, blue indicates synergistic $\kappa$ values, yellow highlights higher IAE50 or IAE90 values, and red highlights lower IAE50 or IAE90 values.

| Cell line | IDMA | IDMB | Kappa ($\kappa$) | $Kappa_{lo}$ | $Kappa_{hi}$ | IAE50 | IAE90 |
|---|---|---|---|---|---|---|---|
| ES-8 *WT* | 289.99 | 0.00 | -0.31 | -0.31 | -0.31 | 23.25 | 18.54 |
| ES-8 *WT* | 303.50 | 0.00 | -0.37 | -0.43 | -0.32 | 22.67 | 18.30 |
| ES-8 *SLFN11*$^{-/-}$ | 246.47 | 0.33 | 0.86 | 0.52 | 1.30 | 1.72 | 1.00 |
| ES-8 *SLFN11*$^{-/-}$ | 248.33 | 0.06 | 0.84 | 0.41 | 1.33 | 3.70 | 1.00 |
